# Library transgenesis in zebrafish through delayed site-specific mosaic integration for in vivo pooled screening of transgenes

**DOI:** 10.64898/2026.01.30.702415

**Authors:** Shahar Bracha, Adam Amsterdam, Yasu Xu, Liyam Chitayat, Anubhav Sinha, Edward Boyden

**Affiliations:** McGovern Institute for Brain Research, MIT, Cambridge, MA; HHMI, Cambridge, MA; Yang Tan Collective, K. Lisa Yang and Hock E. Tan Center for Molecular Therapeutics in Neuroscience, K. Lisa Yang Center for Bionics, MIT, Cambridge, MA; Departments of Brain and Cognitive Sciences, Media Arts and Sciences, and Biological Engineering, MIT, Cambridge, MA; Koch Institute For Integrative Cancer Research, MIT, Cambridge, MA

**Author notes:** Contributed by Edward Boyden.

**Keywords:** protein engineering, neurobiology, synthetic biology, transgenesis, technology development

## Abstract

Functional screening through systematic deletion, editing or addition of libraries of genes is a powerful approach for discovering gene functions and developing improved molecular tools. However, due to the need for high throughput, such campaigns are typically conducted in vitro, leading to many discoveries, especially tools and therapeutics, which fail to translate in vivo. Tissue context, cellular physiology, and systemic regulation shape both tool performance and gene function in ways that simplified culture systems cannot predict. Pooled in vivo screening methods have the potential to enable screening within living animals while preserving the physiological context, but current approaches using viral vectors face three critical limitations: multi-transgene insertions per cell confound genotype-phenotype association, viral packaging constrains transgene size, and cell-type tropism restricts and biases targeting. Here, we introduce a zebrafish library transgenesis method that overcomes these limitations through delayed site-specific mosaic integration. We exploit a temporal delay between library microinjection with PhiC31 mRNA, and library integration, to allows the library to spread episomally throughout the developing embryo before integration begins. This produces mosaic animals where each cell independently integrates one randomly-selected library member, enforced by a single genomic AttP landing site. We demonstrate delivery of multi-kilobase transgenes with high library coverage of 1,378-1,989 unique integrants per animal, and single-transgene-per-cell in ∼99% of brain cells. This method provides a platform for direct in vivo screening of large transgene libraries with single-transgene precision, with potential applications in both biological discovery and tool development.

## Introduction

Systematic screening of genetic libraries, which involves testing many perturbations or transgenes in parallel, can critically accelerate the development of molecular tools and therapeutic interventions. Library screening must balance two competing demands: throughput (the number of variants that can be tested simultaneously), and predictive accuracy (how faithfully screening conditions represent the biological context where the gene products will ultimately function). In vitro screening is typically employed due to the accessibility of high throughput assays, but it often involves sacrificing the predictive accuracy of a screen in critical ways, as in vitro conditions fail to recapitulate the complex cellular and physiological environments that govern gene function in vivo (1–6). Evidently, many genetically-encoded tools developed in vitro have been later found to have diminished or no activity when applied in vivo. For example, genetically-encoded tools can undergo altered processing and trafficking in vivo, including aggregation or mislocalization to unintended subcellular compartments and tissues, which does not manifest during in vitro testing, as has been the case for many voltage indicators (7, 8) and soma-targeted biosensors (9, 10). Tools optimized in vitro have been found to exhibit unexpected off-target effects, non-specificity and deleterious interactions with endogenous processes that are present in vivo but absent in vitro. Such has been the case for early calcium indicators (11, 12), bioluminescent proteins (13), Cas9 (14) and base editors (15). These side-effects can even manifest as toxicity or immunogenicity in vivo, sometimes leading to clearance of the tool or death of the expressing cells, issues which have been faced for Cas9 (16), Cre recombinase (17), early calcium indicators (12, 18) and some optogenetic tools (19, 20). These context-dependent differences mean that many candidates identified through in vitro screening fail during subsequent in vivo validation, wasting time and resources despite extensive optimization efforts (4, 6).

Screening directly in living animals would preserve the physiological context necessary for accurate prediction of in vivo performance. However, traditional approaches for in vivo testing involve expressing and phenotyping one transgene in each animal. This one-by-one testing is prohibitively slow, expensive, and labor-intensive, in addition to requiring special ethical consideration when screening libraries containing hundreds of variants (2, 3). Researchers have thus been limited to either high-throughput in vitro screening, which offers efficiency at the cost of predictive accuracy, or low-throughput in vivo testing that accurately represents physiological context but is impractical for screening at scale. To address this tradeoff, new techniques have been emerging to enable direct in vivo pooled screening - multiplexed testing of many genetic perturbations or transgenes within single animals. This is achieved using transgenesis and mutagenesis methods that deliver pools of genetic modifications to a single animal. These methods create mosaic animals, where different cells in the body harbor different genetic modifications. In this way, many variants can be tested simultaneously in each animal, preserving the in vivo physiological context while enabling higher throughput than one-by-one transgenic approaches (21).

Current pooled in vivo screening methods have focused on perturbation screens, using libraries of gRNAs or siRNAs delivered via viral vectors in rodents (1, 22–24). These methods have been applied to study the effects of endogenous genes within different cell types in vivo, by associating their perturbation (by knockout, knockdown, or mutation) with readouts such as cell survival and proliferation (measured through enrichment or depletion of gRNA/barcode counts in bulk tissue sequencing), gene expression profiling (via single-cell RNA-seq) (1, 22–26), and more recently, imaging-based readouts on fixed tissue (via in situ antibody staining or fluorescence in situ hybridization) (27). While these approaches have been powerful for understanding endogenous gene function in diverse cell types within their native in vivo context, they face three fundamental limitations. First, because transgenesis via viral infection follows a Poisson distribution, there is an unavoidable tradeoff between the number of transduced cells and the proportion of cells transduced with multiple transgenes (24, 25). Cells harboring multiple transgenes confound interpretation by mixing the effects of different genetic modifications, introducing artifacts that are difficult to correct and reducing the effective throughput of the screen. For example, in low-titer lentiviral delivery of gRNA libraries to the cortex of mouse embryos, even sparse targeting of <0.1% of cells resulted in 46% of transduced cells containing multiple perturbations (22). Follow-up studies using optimized AAVs with enzymatic integration led to similar proportions of multi-transgene cells at 2% tissue targeting (24). Second, viral vectors exhibit cell-type-specific tropism that restricts which cell types and tissues can be targeted with sufficient throughput for screening, and biases which cells are targeted for gene phenotyping (28). Viral tropism can also depend on cellular state and other factors that may vary unpredictably, introducing further uncontrolled biases into screening results (29–31). Third, viral packaging constraints limit the size of transgenes that can be screened. AAV vectors, the most commonly used viruses for in vivo gene delivery, can package ∼4.7 kb, preventing their use for screening libraries of large protein-coding transgenes and restricting applications to small transgenes. As a result, to our knowledge, all pooled in vivo screens performed to date have been of small interfering RNAs or gRNA libraries (1, 5, 22–24, 26, 32–36).

A different approach for library transgenesis, implemented in *C. elegans*, is TARDIS (Transgenic Arrays Resulting in Diversity of Integrated Sequences) (37). This approach involves injecting pooled libraries into the *C. elegans* gonads and exploiting delayed transgene integration over multiple generations to generate multiple stable transgenic lines. However, TARDIS produces single-transgene animals rather than mosaic animals, and requires raising many animals and characterizing them one-by-one (37). While useful for screening in *C. elegans*, it relies on the nematode’s idiosyncratic assembly of injected transgenes into heritable extrachromosomal arrays, and requires multiple generations, preventing its generalization to other model organisms. Lastly, the reliance on Cas9-mediated chromosomal breaks and homology-directed repair leads to low efficiency transgene integration, especially for constructs larger than 1-2 kb (38, 39), requiring the use of selectable markers and processing many animals (e.g. heat-shocking and selecting the progeny of 1200 worms for a library of 12 promoters in the TARDIS paper (40)).

In this work, we introduce a method for pooled library transgenesis that addresses these limitations, implemented in zebrafish using developmentally delayed site-specific integration. We inject 1-cell embryos containing a genomic AttP landing site with a mixed library of AttB-containing plasmids alongside PhiC31 integrase mRNA. The temporal delay between mRNA injection and integrase translation and maturation allows embryos to complete many rounds of cell division before the integrase become active. During this delay, the injected plasmid library spreads passively throughout the developing embryo, distributing to all tissues regardless of cell type (addressing the tropism limitation of viral methods). When integration begins, it occurs independently in many cells, with each cell integrating only one randomly selected plasmid from its local episomal pool, enforced by the single genomic AttP landing site (addressing the multi-transgene problem). Unintegrated episomal plasmids are subsequently lost through dilution and degradation, becoming undetectable by 3-4 days post-injection (41–43). Because the plasmids are delivered by direct microinjection rather than viral packaging, the method accommodates large multi-kilobase transgenes (addressing the size limitation). This platform for mosaic transgenesis could accelerate screening campaigns of transgenes and genetic perturbation libraries in vivo, while reducing animal use and associated time and labor for direct in vivo screening.

## Results

### Conceptual design for a library mosaic transgenesis method based on delayed site-specific integration with PhiC31

The utility of a transgenesis method for in vivo pooled screening depends on three critical parameters: (1) efficiency (the fraction of cells expressing a transgene), (2) diversity (the number of different transgenes expressed per animal and the distribution of their abundance), and (3) transgene mutual exclusivity (proportion of cells expressing a single transgene). The third parameter is particularly crucial, as cells expressing multiple transgenes create ambiguity about the mapping between transgenes and their phenotype, which confounds screening results.

A common transgenesis method in zebrafish involves injection of the 1-cell embryo with a plasmid containing transposon arms and an mRNA encoding for Tol2 transposase, leading to random multi-copy genomic integration (44). In theory, this method can be used to deliver multiple transgenes per animal, but the random multi-copy integration mechanism of Tol2 means that most cells will express multiple transgenes, violating the transgene mutual-exclusivity required to support in vivo pooled screening. A recently developed method introduced construct integration using the site-specific bacteriophage integrase PhiC31, instead of the transposase, for single-copy genomic integration (45, 46) (**Fig. 1A**). This method was developed to provide a streamlined way to introduce transgenes into validated safe harbor loci, reducing experimental variability imposed by genomic position effects. For this purpose, two safe harbor zebrafish lines were developed by the Mosimann lab, named pIGLET14a and pIGLET24b, which have a single AttP site on chromosome 14 or 24, respectively (46).

**Figure 1:**
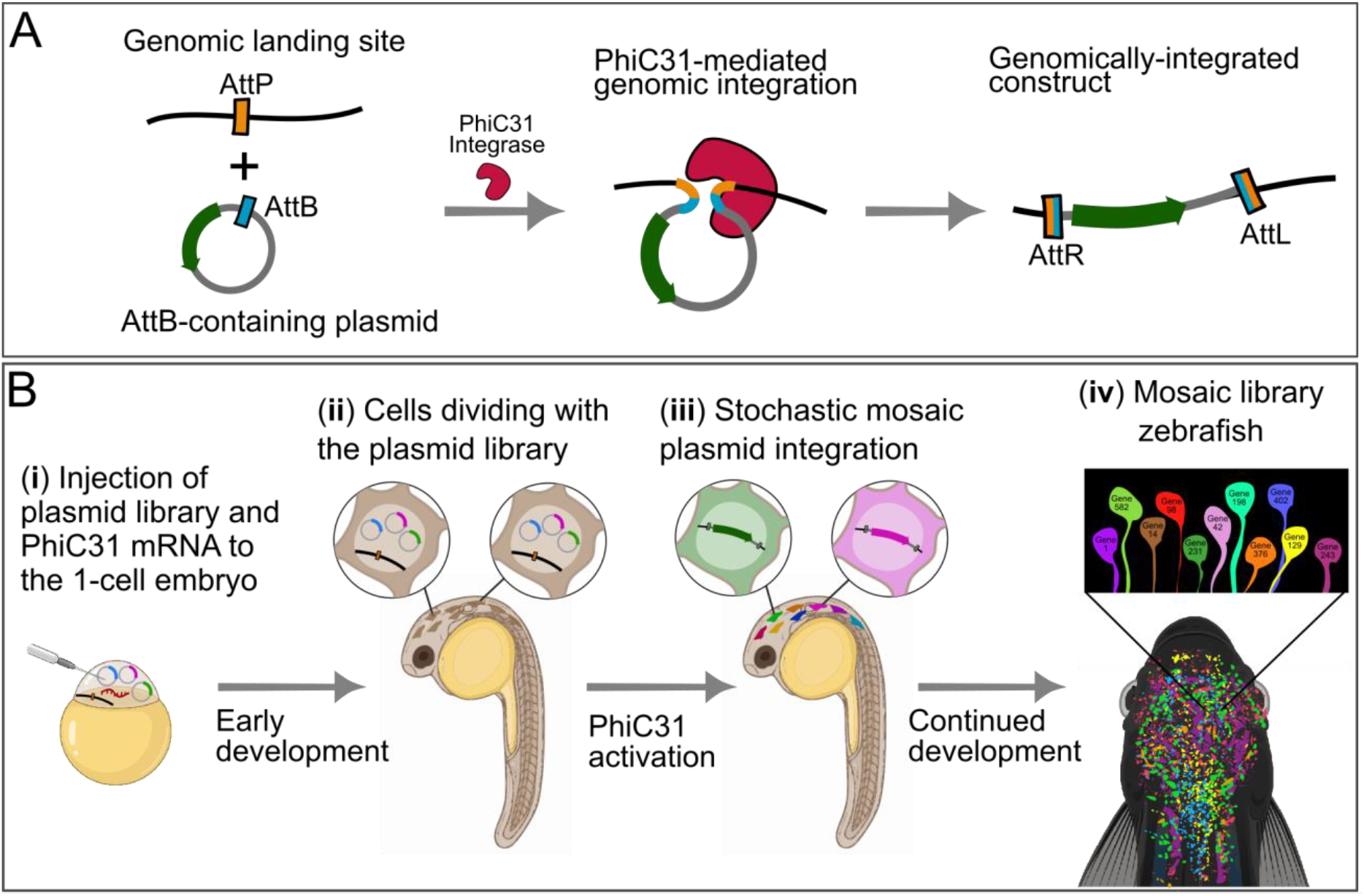
Illustration of the mosaic library transgenesis method. **(A)** A single-copy genomic AttP landing site (orange) in the pIGLET (‘phiC31 Integrase Genomic Loci Engineered for Transgenesis’) line (46), with an exogenously introduced plasmid containing an AttB site (blue) and a transgene cassette (green). Once the PhiC31 integrase enzyme (red) is introduced, it catalyzes recombination between the AttP and AttB, leading to single-copy genomic integration of the plasmid. **(B)** Schematic of the overall procedure of delayed site-specific library transgenesis. (i) The 1-cell embryo is injected with a mixture of plasmids (the transgene library, drawn as circles with blue, magenta and green rectangles) and mRNA encoding for the PhiC31 integrase (red). (ii) During early development, the library passively spreads in the embryo as episomal plasmids together with the PhiC31 mRNA/protein as the cells divide. (iii) After an initial stage of development, the PhiC31 becomes active and integrates a single randomly-selected plasmid from the library in each cell. (iv) This produces a mosaic animal in which different cells express different library members, and only one library member in each cell.

We reasoned that delivery of the integrase via injection of the PhiC31 mRNA, which was part of the original Lalonde et al. protocol (46), could provide a mechanism for enforcing a temporal delay between DNA introduction and integration. Furthermore, we hypothesized that this temporal delay could lead to mutually-exclusive mosaic transgenesis, if a library of plasmids (instead of a single plasmid) was injected (**Fig. 1B(i)**). The exact kinetics of PhiC31 production, maturation and enzymatic activity are unknown, but we hypothesized that multiple embryonic cell divisions could happen before it reached levels sufficient for catalyzing plasmid integration (**Fig. 1B(ii)**). We reasoned that if this is the case, then by the time PhiC31 is active, it could induce many parallel and independent integration events in many different cells (**Fig. 1B(iii)**), resulting in a mosaic animal in which different cells integrated different plasmids from the injected library (**Fig. 1B(iv)**). Importantly, we reasoned that the presence of only one genomic AttP landing site will enforce that each cell integrates only one construct, in a mutually-exclusive manner (37, 47). This is because when the integrase catalyzes recombination between the single genomic AttP site and a plasmid-borne AttB site, it replaces the genomic AttP with AttL and AttR sites that cannot serve as substrates for further integration (**Fig. 1A**).

### PhiC31-mediated mosaic integration leads to high brain targeting with mutually-exclusive transgene expression

To test the transgene mutual-exclusivity of the proposed mosaic transgenesis method, we injected heterozygous pIGLET zebrafish embryos with a simplified two-member library of transgenes. The library consisted of a green and a red fluorescent protein (GFP and mScarlet), each with a membrane tag. The transgenes were expressed pan-neuronally using a non-repetitive UAS (4xnrUAS) promoter in a HuC::Gal4 driver line (48). We injected this 50:50 mixture, together with PhiC31 mRNA, into 1-cell embryos from a cross of pIGLET14a or pIGLET24b females (carrying a single genomic AttP landing site) and HuC::Gal4;nacre;RH1::DsRed males (providing pan-neuronal Gal4 expression required to activate the 4xnrUAS minimal promoter in the integrated constructs). We reasoned that if construct integration is truly mutually-exclusive, the vast majority of fluorescent neurons will be either green or red, and not both. If integration was not mutually exclusive, we would expect to see many cells which are both green and red.

3 and 5 days post-fertilization (dpf), we imaged the larvae to assess the distribution of fluorescent neurons. This allowed us to estimate both the total number of fluorescent neurons (indicating the number of neurons that integrated any construct) and the ratio of single-transgene vs. multi-transgene neurons. We counted fluorescent neurons from eight mosaic animals total. For six of the animals, we counted 1-5 representative FOVs for each (2,473 neurons total), and for two animals we counted neurons across the entire hindbrain volume (2,511 neurons total) to get an estimate of total integration levels in the brain.

Our analysis showed that 99.34% of neurons expressed either GFP or mScarlet exclusively, while only 0.66% (33/4,984 neurons) expressed both fluorophores (**Fig. 2, Table S1)**. The 33 double-positive neurons were distributed amongst the eight fish and different brain areas (**Table S1)**. Given that the original library consisted of an equal mix of GFP and mScarlet, we reasoned that the probability of integrating both at least one GFP and at least one mScarlet construct must be similar to or larger than the sum of the probabilities of integrating multiple GFP-only constructs or multiple mScarlet-only constructs (which would also appear as GFP-only or red-only fluorescence). Based on this assumption, we estimate the total ratio of neurons that integrated multiple transgenes to be up to double the ratio of red-and-green neurons, translating to up to around 1.3% multi-transgene neurons. We hypothesize the rare double-positive cells to be due to spontaneous genomic integration of naked plasmid DNA, a phenomenon known to occur at low frequency without enzymatic mediation, with consistent rates reported in the literature (41). This suggests that PhiC31 mosaic integration is indeed mutually exclusive, and the frequency of multi-transgene cells achieved with this method is sufficiently low for high quality in vivo pooled screening.

**Figure 2:**
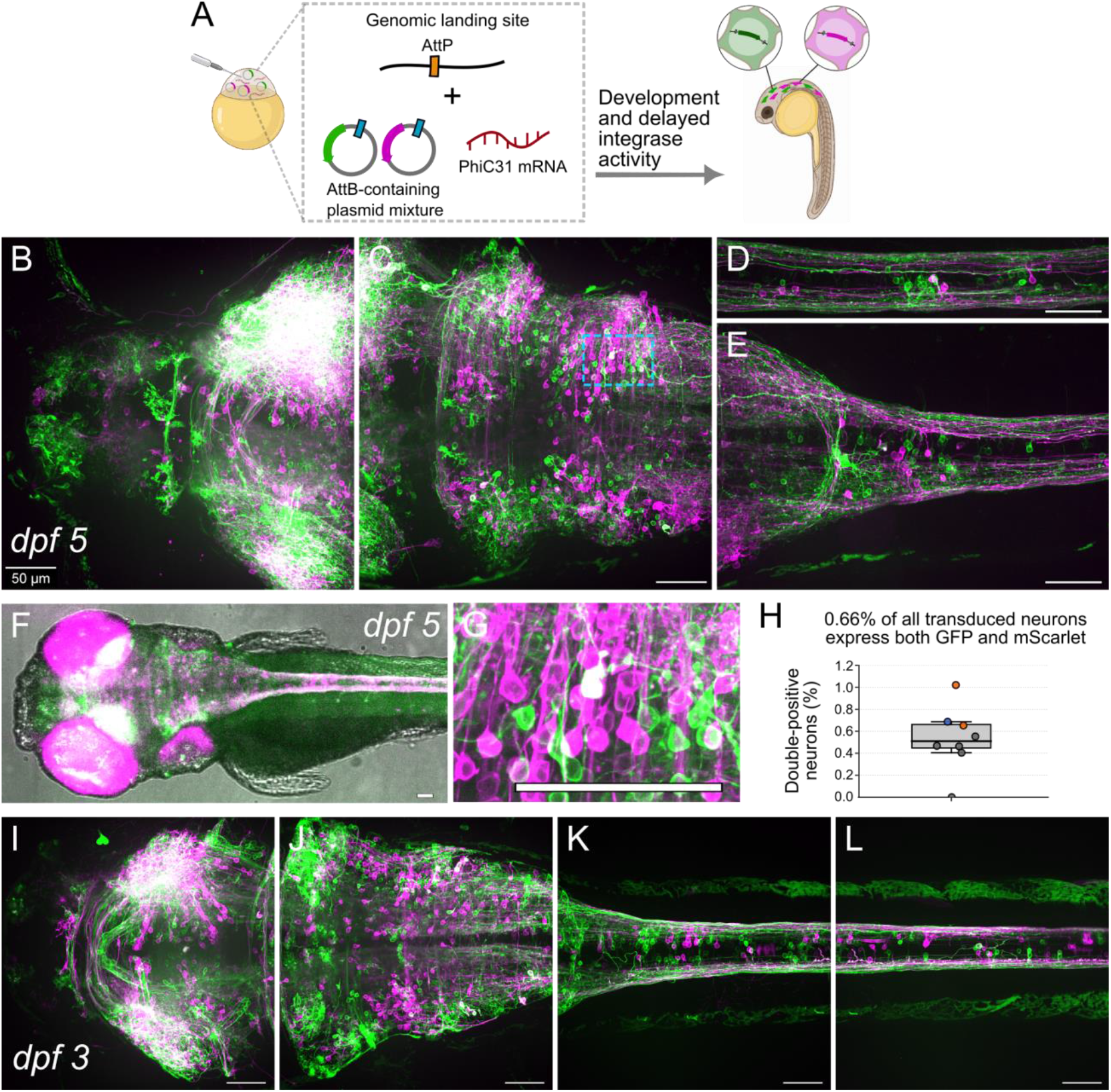
Mutually exclusive expression of library transgenes. pIGLET heterozygous larvae display ∼99% mutually exclusive mosaic expression of a single library member per neuron. (**A**) Illustration of the experiment: pIGLET heterozygous embryos containing a single AttP site in chromosome 14 or 24 were injected with PhiC31 mRNA and a 50:50 mixture of plasmids containing an AttB site (blue) and constructs for neuronal expression of mScarlet (magenta) or GFP (green), fused to a CAAX tag for membrane targeting. 3 or 5 days later, the larvae were imaged. (**B-E**) Representative images of a mosaic 5 dpf pIGLET24b heterozygous animal (pIGLET24b;HuC::Gal4;nacre;RH1::DsRed) following library transgenesis, showing the forebrain and midbrain (B), midbrain and hindbrain (C), posterior hindbrain and spinal cord (D) and spinal cord (E). Max-projection images are shown with skin autofluorescence removed to aid visualization. (**F**) One plane of the fluorescent channels overlaid on a brightfield image of a mosaic 5 dpf pIGLET14a heterozygous larva (pIGLET14a;HuC::Gal4;nacre;RH1::DsRed, with prominent expression of the red eye marker). (**G**) Zoomed-in image of the section marked with a cyan box in the hindbrain in (C), showing a neuron co-expressing GFP and mScarlet. (**H**) Quantification of the ratio of neurons expressing both GFP and mScarlet (double-positives), out of all transduced neurons, in 8 mosaic larvae. The animals for which the whole hindbrain was quantified are marked in orange. The 3 dpf larva is marked in blue. The rest (marked in gray) are 5 dpf larvae for which 1-4 random FOVs were quantified. All the raw numbers are available in Table S1. (**I-L**) As in B-E, but for a 3 dpf larva. Scale bar: 50 μm.

We estimated the total frequency of targeted neurons across the brain based on the counts of fluorescent neurons across the entire hindbrain volume of two representative 5 dpf larvae, which amounted to an average of 1,255 fluorescent neurons per hindbrain (980 in Fish 7 and 1,531 in Fish 8) (**Table S1)**. The hindbrain was chosen as it enabled the most accurate identification of fluorescent neuronal somata, thanks to its structure and distance from the eyes. In more detail: the HuC::Gal4 driver line used contains a multi-copy red fluorescent eye marker (RH1::DsRed) as a genotyping aid, which exhibits variable expression levels. Therefore, in some fish, brain areas close to the eyes (the optic tectum and forebrain) exhibited a red fluorescent glow, under confocal microscopy, which interfered with the identification of mScarlet-expressing neurons in those areas. In addition, the high density of axons in the optic tectum around the eyes interfered with the identification of neuronal somata (both green and red) in those areas. We used the published Z brain atlas (zebrafishexplorer.zib.de, (49)) and dataset from Ahrens at al (50, 51) to estimate the total number of neurons in the hindbrain of 5 dpf larvae to be around 25k. Given this estimate, our quantification of around 1,255 fluorescent neurons across the hindbrain suggests that approximately 5% of neurons expressed library integrants, assuming that the rate of integration was similar in all areas of the brain. The latter was consistent with our qualitative assessment of 25 imaged fish over 6 independent experiments (**Fig. 2)**. Those fish showed qualitatively similar levels and distribution of fluorescent neurons across the brain.

### Quantifying the number and distribution of library transgenes integrated in mosaic animals

After establishing the overall efficiency and mutual exclusivity of library integration, we set out to quantify the maximum library diversity achievable per animal. We reasoned that the total number of different transgenes that can be expressed in one animal will be determined by the number of independent integration events, which is affected by the developmental stage at which PhiC31 becomes active. Prior studies following embryos after injection of mRNA encoding for GFP showed that significant green fluorescence is detectable at 3 hours post-fertilization (hpf) (52). Consistent with that, evidence shows that after injection of PhiC31 mRNA to the 1-cell embryo, construct recombination can be observed as soon as 3.3 hpf (45). By 3.3 hpf, a zebrafish embryo is estimated to contain 1-2k cells (53). Of course, given the likely possibility that integration events are distributed over time, the actual number of independent integration events could be lower if most of them occur before 3.3 hpf, or higher if they continue occurring at later stages as well. It also depends on the dynamics of retainment of the PhiC31 mRNA/protein and of the episomal library of AttB-containing plasmids as cells divide. The latter likely depends on multiple factors, including: (i) how many plasmid and mRNA molecules were originally injected, (ii) the rate of degradation of the episomal plasmids, mRNA and PhiC31 protein, and (iii) the uniformity of the distribution of the episomal plasmids and integrase mRNA/protein among the embryo cells as they divide.

Therefore, we set out to empirically quantify the number of independent integration events that occur in our protocol. We generated a library of plasmids containing 15-nucleotide random DNA barcodes preceding a GFP-CAAX expression cassette (**Fig. 3A**). This design allowed us to use bulk deep sequencing to quantify the diversity of integrated transgenes based on the DNA barcodes recovered from each mosaic animal. We injected the barcoded library into 1-cell heterozygous pIGLET embryos (pIGLET24b;HuC::Gal4;nacre;RH1::DsRed) and imaged larvae with widespread neuronal GFP expression at dpf 5 (**Fig. 3B**). After imaging, we extracted genomic DNA from their entire body and amplified the pool of integrated barcodes using 10-cycle PCR with primers spanning the genomic integration junction. The primers were designed to ensure that only barcodes from transgene constructs that integrated into the AttP landing site got amplified, and to exclude the possibility of amplifying barcodes from unintegrated episomal plasmids. This was achieved using a forward primer targeting a genomic sequence upstream of the AttP on chromosome 24, and a reverse primer targeting the plasmid-specific HS4 insulator sequence downstream of the barcode location. This resulted in a 650 bp amplicon library containing the integrated barcodes from each larva (**Fig. 3C**). The amplicon libraries from 12 imaged larvae that displayed high levels of neuronal GFP expression and strong amplicon bands (**Fig. S2**) were further processed to generate sequencing libraries. The 650 bp amplicon library from these larvae were re-amplified with a 15-cycle PCR, using primers that were internal to the first ones, and added illumina-compatible sequencing overhangs and sample-specific 5-nt barcode to the amplicons from each larva, to allow for multiplexed pooled sequencing. Deep sequencing yielded a total of 7.2 million high-quality reads across the 12 fish samples. In addition, we sequenced the source plasmid library by amplifying the barcodes from the original injected plasmid sample, which yielded 5.8 million sequencing reads (**Table S2)**. The 15-nt barcode was extracted from each read based on a sequence search for the conserved sequences surrounding the barcode. We collapsed all closely related barcodes (defined as Levenshtein distance of 1 apart), reasoning that given the high complexity of the original library, the likelihood of such closely-related barcodes appearing in sequenced fish by chance is far exceeded by the likelihood of 1-nt sequence divergence resulting from sequencing errors or mutations introduced during amplification (**Table S3, Fig. S4**). The fact that most clusters of closely-related barcodes consisted of one high-count barcode and multiple low-count barcodes further support this assumption. We filtered out rare barcodes that appeared less than 3 times in a given fish, to further exclude rare barcodes that could represent potential sequencing errors, contaminants, or errors in the barcode extraction. While the proportion of reads with rare barcodes was low in the fish samples (<1% of reads, **Table S3**), in the source barcode library, most of the reads contained rare barcodes (appearing <3 times in the sequenced library), suggesting that the complexity of the injected library was substantially higher than the sequencing depth. In accordance with this observation, we used a more permissive threshold for rare barcode inclusion in the source library (removing only barcodes with count<2, which accounted for 45% of the reads). Overall, our analysis revealed that each mosaic animal integrated on average 1,676 different barcodes (median: 1,682, Std. Dev.: 176, range: 1,378-1,989) (**Fig. 3D, Table S3)**. As mentioned above, analyzing the original injected barcode library revealed that it had very high complexity, with no barcode being represented at >0.0005% frequency in the library even after collapsing closely-related barcodes. This makes it likely that for each fish, each unique barcode observed originated from a single integration event.

**Figure 3:**
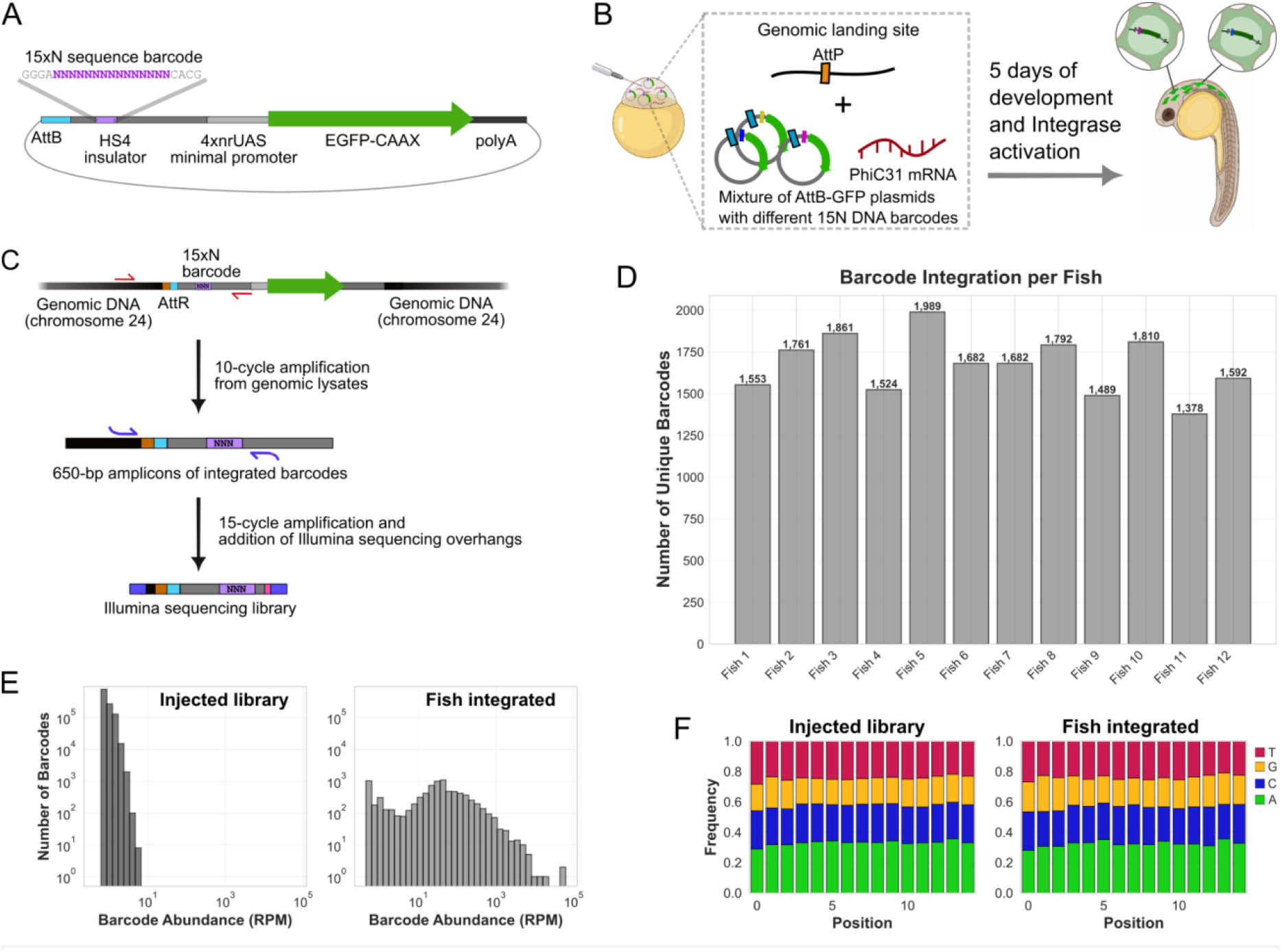
quantifying the capacity of delayed integrase library transgenesis with DNA barcodes. (**A**) Construct design for the barcoded GFP-CAAX library plasmids. Each plasmid includes an AttB sequence (light blue), HS4 insulator element (medium gray) embedded with a unique 15-nt random sequence barcode (15xN, purple), a 4xnrUAS minimal promoter for tissue-specific Gal4 transcription activation (light gray), GFP-CAAX (green arrow) and SV40 polyA (dark gray). (**B**) Scheme illustrating the experiment-a high-complexity library of barcoded GFP-CAAX plasmids, each containing a different 15xN barcode (dark blue, magenta and yellow rectangles on the plasmids), was injected into 1-cell embryos of heterozygous pIGLET24 zebrafish containing a genomic AttP site on chromosome 24. The library was co-injected together with mRNA encoding for the PhiC31 integrase (red). 5 days later, the larvae were imaged to confirm GFP expression in neurons, and then selected for extraction of genomic DNA from their entire bodies (n=12 larvae). (**C**) the genomic extracts were used as templates for 10-cycle PCR amplification of the barcodes from all genomically integrated plasmids, using primers that specifically target the integration junction (shown in red arrows). The resulting 650-bp amplicon library is then re-amplified with different, internal primers (shown in blue arrows), to attach overhangs for Illumina next generation sequencing (blue) and add a 5-nt sample-specific multiplexing barcode (pink) for pooled sequencing. (**D**) Number of unique high-confidence barcodes identified in each fish, after barcode collapsing and filtering. (**E**) Histogram of barcode abundance for the injected source library and for the barcodes recovered from the fish-integrated plasmids. The injected library displays a narrow distribution, indicating high complexity (many rare barcodes appearing at similar low frequency) while the fish-recovered barcodes display a broader long-tailed distribution (some barcodes appearing much more than others), consistent with intra-fish clonal expansion of the integrated transgenes. The per-fish barcode abundance histograms are available in Fig. S5. RPM=reads per millions (read counts normalized to the total number of reads sequenced for each sample). (**F**) Nucleotide composition for each position in the injected library and in the integrated barcodes shows high sequence diversity and no sequence bias for integration. Per-fish barcode sequence compositions are available in Fig. S2.

Barcode sequence analysis revealed no favored sequence composition or motifs for the barcodes from the integrated plasmids compared to the injected library (**Fig. 3F, Fig. S3**). Furthermore, the sequence diversity, measured by the average pairwise hamming distance between the barcodes integrated in each fish was the same as that of the injected library (11.1), and similar to the expected value for a theoretical uniform random sequence library (11.3), indicating that the different barcodes identified in each fish are indeed random and are the products of independent integration events, rather than from barcode diversification by mutation of a small number of integrants in each fish (**Fig. S4**).

When looking at the distribution of barcode representation, we found clear signs of clonal expansion of the barcodes within the mosaic animals, as expected. While the sequenced source library displayed millions of different barcodes, each represented at similar rare frequency, the mosaic animals tended to have a different set of around 1,600 barcodes each, and the barcodes were represented at different frequencies (**Fig. 3E, Fig. S5, Fig. S6**). As the high library complexity suggests that each unique barcode originated from a single integration event, variance in barcode abundance is most likely the result of clonal amplification as cells replicated after barcode integration, rather than multiple independent integration events of the same barcode. After a cell integrates a library member, all its progenies will inherit the same barcode, creating clonal populations whose size reflects the number of cell divisions between integration and gDNA extraction at 5 dpf. Therefore, the variance in integrated barcode abundance could represent either (1) variable integration timing - barcodes integrating earlier in development undergo more amplification with cell division; or (2) variable proliferative capacity of different cell lineages-namely, barcodes integrating into highly proliferative lineages (e.g. basal stem cells) achieving greater expansion than those in slowly dividing or post-mitotic cell types (e.g. neurons). Accordingly, we would expect that the variance in transgene representation would be lower in library transgenesis applications involving cell type- or tissue-specific expression (for example, when screening transgenes only in neurons). In such applications, the total number of different transgenes expressed per animal would also be smaller, since our DNA barcode quantification included all barcodes integrated across the entire body of the zebrafish.

## Discussion

We developed a method for library transgenesis that provides a platform for high-throughput in vivo screening. By exploiting a temporal delay between plasmid library injection and PhiC31-mediated integration in zebrafish, we achieve mosaic transgenesis with 1,378-1,989 unique integrated transgenes per animal and ∼99% mutually-exclusive transgene expression. Site-specific integration to a single genomic landing site ensures that nearly all transduced cells express a single transgene while still enabling delivery to many cells across the tissue, circumventing the tradeoffs that limit methods relying on stochastic infection and integration events such as viral delivery, transfection and random transposase-mediated genomic integration. By creating mosaic animals in which different cells express different transgenes, we can effectively transform each animal into hundreds of parallel experiments. This could enable high-throughput screening of transgene libraries in the native in vivo physiological context, which would be valuable for developing better genetically-encoded tools through direct in vivo screening, and for basic research investigating the effects of libraries of genetic perturbations in vivo.

We demonstrate mosaic somatic transgenesis, where libraries of transgenes are expressed in different somatic cells following delayed integration, allowing each animal to function as a living library with individual cells testing different variants in the native in vivo context. This approach is well-suited for screening genetic perturbations and transgenes with cell-autonomous effects, where the phenotype of a single transduced cell can be reliably assessed even when surrounded by non-transduced or differentially-transduced neighbors. Applications include screening molecular tools in vivo, such as genetically-encoded biosensors, fluorescent markers, DNA editing enzymes and more. Additionally, mosaic somatic transgenesis could be applied to enable lineage tracing and brainbow-like barcoding strategies for morphological tracing and cell segmentation, including for connectomic mapping (54–57). Beyond this demonstrated application, we hypothesize that mosaic germline transgenesis could also be achievable, where injected animals would generate libraries of progeny animals, each containing a single transgene throughout its entire body, similar to the TARDIS approach (37).

While we demonstrate mosaic integration of 1,378-1,989 unique library variants per animal, future optimization could further increase library complexity and integration efficiency. Protocol refinements may include varying the concentrations and purification methods used for the injected plasmid libraries and mRNA and refining the AttB plasmid design. Using optimized hyperactive PhiC31 instead of the native bacteriophage sequence could provide another strategy to enhance integration efficiency (59). It would be interesting to investigate more deeply the timing of integrase expression and activity, and to explore alternative mechanisms for delayed integration. One option could involve introducing the integrase gene as DNA instead of mRNA. For example, if the integrase was encoded on a co-injected DNA plasmid under a ubiquitous promoter, its transcription would only begin around 3 hours post-fertilization with the start of zygotic transcription after 10 embryonic cell divisions (60), likely resulting in an even longer temporal delay before integration. Alternatively, a tissue-specific promoter could further delay integration until after a specific tissue or cell lineage forms, while an inducible promoter (e.g., heat-shock or drug-inducible) would provide flexible spatiotemporal control of integration initiation (61–63). Future approaches could involve generating transgenic zebrafish lines with delayed or inducible integrase expression cassettes in their genome, further increasing overall efficiency and tissue coverage.

In our current implementation, DNA barcode library analysis demonstrated clonal expansion of the integrated variants, which was expected from the mechanism of delayed integration into different cells across the entire body. We hypothesize that this could result from either variance in integration timing or variance in the proliferative capacity of different cell types receiving different integrants. If the latter is the case, we would expect this variance to be significantly lower when the method is applied to screening library variants expressed in a specific tissue or cell type. In addition, more precise control of the variant abundance distribution could be achieved by keeping the injected library smaller than the number of integration events. The overall library complexity (measured both by the number of unique variants and their distribution) achieved here should be suitable for many screening applications, and further improvements could enhance the method’s throughput and efficiency.

We implemented this approach in zebrafish, which offers unique advantages for many of the applications we discuss. Its natural transparency and small size make it highly amenable to imaging-based phenotypic analysis of transgene and perturbation libraries in live mosaic animals. As a vertebrate model, it recapitulates many aspects of human physiology with demonstrated clinical translatability and a wealth of established disease models (64, 65). Critically, this work was enabled by the existence of AttP landing site lines with validated safe-harbor integration sites that were already established for zebrafish (46). Similar engineered lines with genomic integrase landing sites have also been established for C. elegans, Drosophila, pigs and mice (66–71), providing a foundation for adapting this approach to those model organisms as well. As we increasingly recognize that gene functions depend critically on their interactions within complex in vivo environments, including aspects we may not yet even fully understand, methods that preserve this physiological context while enabling high-throughput screening will be essential for both developing better molecular tools and for basic biological discovery.

## Acknowledgements

We thank Christian Mosimann and his lab for the pIGLET zebrafish lines. Thanks to all members of the Boyden lab for many fruitful discussions. Fig. 1B, 2A, 3B contain illustrations from biorender.com. ESB acknowledges, for funding, Lisa Yang, HHMI, NIH 1U01NS120820, NIH 1R01MH123977, NIH R01MH122971, and NIH R01DA029639. SB acknowledges funding from the Y. Eva Tan Postdoctoral Fellowship and the Yang Tan Collective at MIT.

## Methods

### Materials and data availability

All the raw imaging data and Illumina sequencing data associated with Fig. 2, 3, S1, S2, S3, S4, S5, S6, and Table S1, S2, S3, are available on DOI https://doi.org/10.5061/dryad.d2547d8h0. All the code used to analyze the sequencing data is available on https://github.com/shaharbr/library_transgenesis. The full sequences of key plasmids and primers used in this study are available in appendix data S1, S2, S3 and S4. The plasmids AttB-HS4-nrUAS-GFP-CAAX and AttB-HS4-nrUAS-mScarlet-CAAX will be made available through Addgene upon publication.

### Zebrafish husbandry and transgenesis

All procedures were done in accordance with government and university guidelines, and approved by the MIT Committee on Animal Care. Heterozygous pIGLET embryos for injection were obtained by crossing homozygous pIGLET14a or pIGLET24b or pIGLET24b;HuC::Gal4;RH1::DsRed;nacre (for the single-copy AttP landing site) with HuC::Gal4;nacre (for HuC-driven pan-neuronal expression of proteins under the minimal 4xnrUAS promoter) adult zebrafish. Homozygous pIGLET14a and pIGLET24b were obtained as a gift from Prof. Chris Moismann’s lab (46). HuC::Gal4;RH1::DsRed;nacre;nacre zebrafish were generated in-house based on HuC::Gal4;RH1::DsRed obtained as a gift from Prof. Herwig Baier’s lab. Homozygous pIGLET14a;HuC::Gal4;RH1::DsRed;nacre and pIGLET24b;HuC::Gal4;RH1::DsRed;nacre lines were generated in-house by crossing the above. 1-cell embryos were microinjected with approximately 1 nanoliter of injection mix containing 50 ng/μL total plasmid DNA and 50 ng/μL PhiC31 mRNA, mixed in a total volume of 5 μl RNAse-free water with 0.1% phenol red as a visual marker for successful injection. Microinjections were performed using pulled glass capillaries. Embryos were raised at 28°C in aquarium makeup water (Instant Ocean solution diluted to 450 microSiemens and adjusted to pH 7.0 with sodium bicarbonate) until their analysis at 3-7 days-post-ferlitization (dpf).

### Plasmid and mRNA purification

All plasmids for microinjection were extracted using the QIAprep Spin Miniprep Kit (Qiagen cat #27106) without the addition of RNAse in the lysis buffer, and eluted in water. The PhiC31 plasmid (Addgene #68310) was used to produce purified PhiC31 mRNA using the mMESSAGE mMACHINE™ T7 Transcription Kit (Thermo Fisher, AM1344) with lithium chloride purification. Stocks were diluted to 100 ng/μl, aliquoted to 3 μl per tube and kept in the -80C freezer until the experiment. mRNA aliquots were thawed and kept on ice for each experiment, with up to one freeze-thaw cycle per aliquot.

### Generation of the barcoded plasmid library

To generate the DNA barcode library (barcoded AttB-HS4-15N-nrUAS-GFP-CAAX), we added 15-nt random sequence barcodes into a base plasmid encoding for expression of membrane-targeted GFP (AttB-HS4-nrUAS-GFP-CAAX). The base plasmid contained an AttB site followed by a HS4 insulator sequence, 4xnrUAS (non-repetitive UAS) minimal promoter (48), GFP fused to CAAX membrane targeting motif and a polyA sequence, in a pTwist backbone.The GFP expression was used as validation to confirm successful injection and integration in the larvae that were picked for genome extraction and sequencing. The 15-nt random sequence DNA barcodes were added in the middle of the HS4 insulator sequence using 2-cycle PCR with a reverse primer containing 15 random bases (N’s), followed by a NheI restriction site and extra homology handles for further amplification (primers *F_HS4_NheI* and *R_Add15N_HS4_NheI* in Appendix data S4). The 2-cycle PCR was performed on 20 fmol plasmid (∼50 ng) with 400 fmol of each of the primers (20-fold molar excess), with Q5 high fidelity polymerase mix (cat #M0492 NEB). The reactions were incubated in a thermocycler with initial denaturation (98C, 30 sec) followed by 2 cycles of: 98C for 10 sec, 69C for 30 sec, 72C for 120 sec, and then final extension (72C, 2 min). The products were incubated with 1 μl of the restriction enzyme DpnI for 2 hours at 37C (to eliminate the template plasmid), followed by heat inactivation for 20 min at 80C, and column extraction. The purified products were then re-amplified with primers homologous to the ends of the 2-cycle PCR primers (primers *F_amp_HS4* and *R_amp_HS4* in Appendix data S4). PCR was performed with Q5 high fidelity polymerase mix, with initial denaturation (98C, 30 sec) followed by 30 cycles of: 98C for 10 sec, 67C for 30 sec, 72C for 120 sec, and then final extension (72C, 2 min). The products were then incubated for 2 hours at 37C with 1 μl each of the restriction enzymes DpnI and NheI (to expose the sticky-ends at the ends of the amplicons), followed by heat inactivation for 20 min at 80C. Then, the products were gel extracted and eluted in 20 μl water. Half (10 μl) of the eluted product was then circularized by ligation with T4 ligase (cat #M0202 NEB) for 15 min in room temperature, followed by heat inactivation at 65C for 10 minutes. 5 μl from the resulting ligation product was transformed into e. coli as a pooled library. 1% of the transformed bacteria were plated onto an agar plate with 100µg/mL carbenicillin, and the remaining 99% was grown in a 40 ml liquid culture of LB with 100µg/mL carbenicillin overnight. The liquid cultures were used to midi-prep the library using the QIAGEN Plasmid Plus Midi Kit (Qiagen cat #12943) without the addition of RNAse in the lysis buffer, and eluted in water. Successful cloning and library complexity quality controls were estimated by individual whole-plasmid sequencing of 5 random colonies from the plated transformed bacteria, and by nanopore sequencing of 10,000 plasmid reads.

### Imaging and analysis of the mosaic larvae expressing fluorescent protein libraries

The images shown in **Fig. 2** and **Fig. S2** were acquired of 3-5 dpf mosaic larvae, which were injected with a 50:50 mixture of plasmids for pan-neuronal expression of GFP-CAAX or mScalet-CAAX (AttB-HS4-nrUAS-GFP-CAAX and AttB-HS4-nrUAS-mScarlet-CAAX) as 1-cell embryos.

The images shown in **Fig. S1** were acquired of 5 dpf mosaic larvae injected with the plasmid library of barcoded GFP-CAAX (barcoded AttB-HS4-15N-nrUAS-GFP-CAAX) as 1-cell embryos.

3 dpf larvae were imaged live, while 5 dpf larvae were fixed in 4% PFA overnight, washed three times in aquarium makeup water and mounted in 1% low-melting agarose. Mounted fish were imaged using a spinning disk confocal microscope (Yokogawa CSU-W1 Confocal Scanner Unit on a Nikon Eclipse Ti microscope) with 10x air objective and a 40x water immersion objective (Nikon MRD77410). The microscope is equipped with a Zyla PLUS 4.2 Megapixel camera controlled by NIS-Elements AR software, and laser/filter sets for 405 nm, 488 nm, 561 nm and 640 nm optical channels. We acquired 2-4 FOV for each fish, each as a z-stack with 2.5 μm intervals, to cover most of the brain volume. The images shown in **Fig. 2B-F and Fig. S1** are representative of results obtained over six repeats of the experiment, with 25 larvae total imaged aged 3-7 dpf. The images shown in **Fig. 2, S1 and S2** are max-intensity projections, generated using the ImageJ Z Project plugin. For **Fig. 2 and S1**, green autofluorescence from the skin was removed using manually-drawn masks on the individual z-planes, before generating the max-intensity projection, as demonstrated in **Fig. S2**. This was done to prevent the obstruction of neurons in one plane by skin autofluorescence in adjacent planes, which would otherwise cover and hide them in the max-projection. All the raw imaging data (.nd2 hyperstacks) associated with **Fig. 2, S1, S2** and **Table S1**, are available on are available on https://doi.org/10.5061/dryad.d2547d8h0. The cell counting quantification in **Table S1** was performed on eight larvae aged 3 or 5 dpf, from 6 different injection clutches over 3 independent experiments. Counting of red and green fluorescent neurons was performed manually using the ImageJ CellCounter plugin.

### DNA extraction from zebrafish

For the results shown in **Fig. 3D-F, Fig. S1, S3, S4, S5, S6** and **Tables S2 and S3**, zebrafish larvae were evaluated for GFP expression at 5 dpf, and after confirmation of GFP expression (**Fig. S1**), 24 were picked for euthanization and lysis for genomic extraction. The larvae were lysed in 180 μl Qiagen buffer ATL (cat #19076) with 20 μl proteinase K (20 mg/mL, cat #19134) and incubated in 56C for 1 hour with vortexing every 20 minutes, followed by 90C for 20 minutes. The lysed samples were then processed with the QIAamp DNA FFPE Tissue Kit (cat #56404) using the manufacturer’s protocol, starting from the lysis section. The final genomic DNA was eluted in 30 μl water.

### Amplicon library generation from zebrafish amplified integrated barcodes and from the source injected library

The genomic extracts from 24 larvae were amplified to produce a 652 bp amplicon of the integrated plasmids (illustrated in **Fig. 3C**). These amplicons were generated by PCR with a forward primer on a genomic location on chromosome 24, ∼340 bp upstream of the AttP landing site (*F_Chr24pIGLET*) and a reverse primer on the plasmid in the HS4, ∼200 bp after the barcode (*R_HS4*), for 10 cycles, with Q5 high fidelity polymerase, using the following conditions: initial annealing with 98C for 30 s, 10 cycles of: 98C for 10 s, 70C for 30 s, 72C for 40 s, and final extension with 72C for 2 min. We used 10 μl (a third) of each genomic extract as template. Then, the PCR products were run on a gel and the 652 bp amplicons were extracted from the gel and eluted in 25 μl water. At this point, we selected 12 of the 24 fish-derived amplicon samples for further processing, based on the density of their amplicon bands on the gel, and based on the high level of neuronal GFP expression recorded in the larvae they originated from (**Fig. S1**). 20 μl of each purified amplicon library was used as template for a second PCR reaction using primers internal to the first, which also added 5-nt sample multiplexing barcodes and Illumina sequencing overhangs in the forward and reverse direction (*F_Chr24_illumread* and *R_HS4_FishX_illumread*). This PCR went for 15 cycles using similar conditions to the above, and after it the 325 bp amplicons were run on a gel and purified as above. The resulting purified amplicons from the different barcoded samples were then combined into one pooled sequencing library (illustrated in **Fig. 3C**). To generate sequencing reads from the source injected plasmid library, we amplified the original plasmid sample with primers that bind upstream and downstream of the 15xN barcode in the HS4 element, while adding Illumina sequencing overhangs (*F_HS4_illumread* and *R_HS4_illumread*). This produced 336 bp amplicons for direct Illumina sequencing, with similar PCR conditions and purification as the above.

### Library prep and Illumina sequencing

Sequencing libraries were constructed from amplicon samples using the Illumina DNA Prep tagmentation kit paired with Illumina Unique Dual Indexes, without the tagmentation steps. Libraries were sequenced by SeqCoast Genomics (Portsmouth, NH) on Illumina NextSeq2000 using a 300-cycle XLEAP-SBS flow cell kit, generating paired-end reads (2x150). To ensure accurate base calling, 1-2% PhiX control DNA was added to each sequencing run. Post-sequencing processing, including sample demultiplexing, trimming, and run metrics analysis, was conducted using the integrated DRAGEN v4.2.7 software on the NextSeq2000 platform. Quality assessment (shown in **Table S2**) was performed at two levels: evaluation of overall run performance to confirm sequencing data integrity, and targeted review of FastQC quality reports for individual samples. Overall, sequencing of the fish-derived amplicons produced 7.2 million paired-end reads total and the source library amplicons produced 5.8 million paired-end reads total, with >96% bases with Phred quality score >= 30. After demultiplexing, each fish-derived sample had 538-643k reads (**Table S3**).

### Demultiplexing and barcode extraction

Raw sequencing reads were processed using a custom Python-based analysis pipeline, available at https://github.com/shaharbr/library_transgenesis. For the 12-fish pooled library, reads were first demultiplexed based on the 5-nt sample barcodes (with tolerance for 1 mismatch) and reverse-complemented to correct for sequencing orientation. The injected barcode library reads required no demultiplexing and were processed in their original orientation. For both samples, the 15-nt variable barcodes were extracted by identifying conserved anchor sequences flanking the barcode region: a 12-nt sequence (AGCCCCCAGGGA, allowing 2 mismatches) upstream and a 5-nt sequence (CACGC, requiring exact match) downstream. The extraction algorithm employed progressive search stringency (exact match → Hamming distance up to 2 → Levenshtein distance up to 2) with position validation to ensure accurate barcode identification. The full results from this analysis are included in **Table S3**.

### Barcode collapsing, error correction and high-confidence barcode filtering

To account for PCR and sequencing errors, barcodes differing by a Levenshtein distance of 1 (single nucleotide substitution, insertion, or deletion) were collapsed into a single parent barcode. The parent barcode was defined as the most abundant sequence. All read counts from child barcodes were aggregated into their respective parent barcodes, preserving per-sample information.

For the injected barcode library, barcodes were retained if they had ≥2 reads. For the integrated barcodes from fish, filtering was performed on a per-fish basis: a barcode was retained in a given fish only if it had ≥3 reads in that fish; otherwise, that fish’s count for that barcode was set to zero. Barcodes with Levenshtein distance ≤2 to any conserved (non-barcode) region of the template read structure were removed to eliminate potential artifacts from faulty barcode extraction. The full results from this analysis are included in **Table S3**.

### Barcode abundance and diversity analysis

Read counts for each barcode were normalized to Reads Per Million (RPM) to account for differences in sequencing depth between samples (histograms shown in **Fig. 3E and Fig. S5**). For each sample, we calculated the coefficient of variation (CV), Shannon diversity index (using log base 2), and quartile ratio (Q3/Q1) as measures of barcode abundance distribution (shown in **Fig. S6**). Sequence composition bias was assessed by calculating positional nucleotide frequencies across all barcodes and comparing library and integrated barcode populations (shown in **Fig. 3F and Fig. S3**). Pairwise Hamming distances were computed on 20,000 randomly sampled barcode pairs to quantify sequence diversity (shown in **Fig. S4**).

## Supplementary Information (SI files)

**Figure S1.**
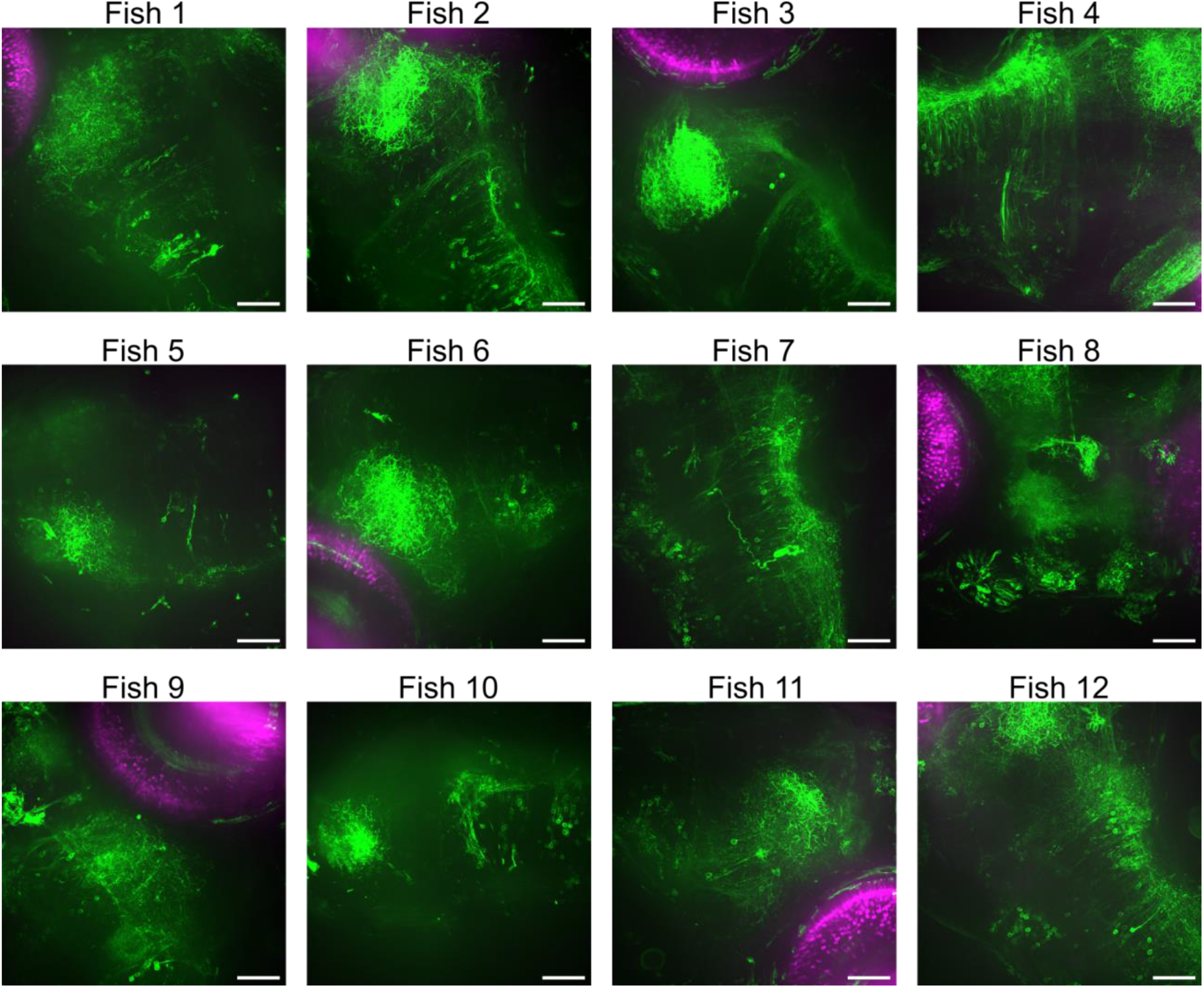
GFP-CAAX expression in the 12 fish with DNA barcodes characterized by deep sequencing (related to **Fig. 3)**. Images presented are max projections from confocal fluorescence imaging of the brain around the optic tectum and/or hindbrain of the animals, with skin autofluorescence removed with manual masks to aid visualization. Magenta fluorescence corresponds to red eye marker expression in the HuC::Gal4;nacre;RH1::DsRed driver fish line. Scale bar = 50 μm.

**Figure S2.**
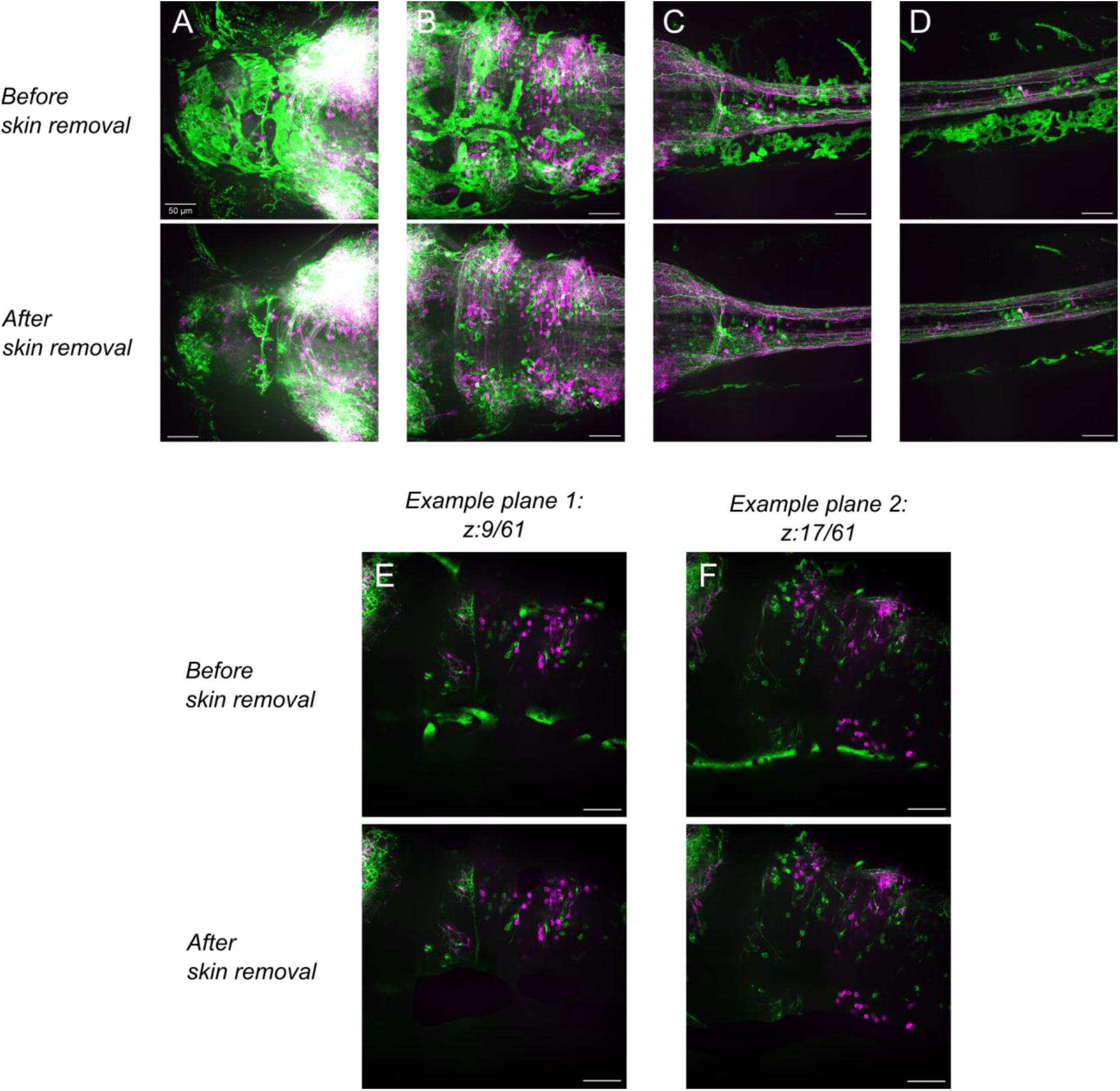
Demonstration of the skin autofluorescence removal from the max-projections images shown in Fig. 2 and S1. Green autofluorescence from the skin was removed using manually-drawn masks on the individual z-planes, before generating the max-intensity projections. This was done to prevent the obstruction of neurons in one plane by skin autofluorescence in adjacent planes, which would otherwise cover and hide them in the max-projection. **Top (A-D)**: max-projections of confocal images from forebrain and midbrain (A), midbrain and hindbrain (B), posterior hindbrain and spinal cord (C) and spinal cord (D), as shown in Fig. 2, before and after removal of the skin autofluorescence. **Bottom (E-F)**: Two examples of individual z-planes from the max-projection shown in (B), before and after removal of the skin autofluorescence. Scale bar = 50 μm.

**Figure S3.**
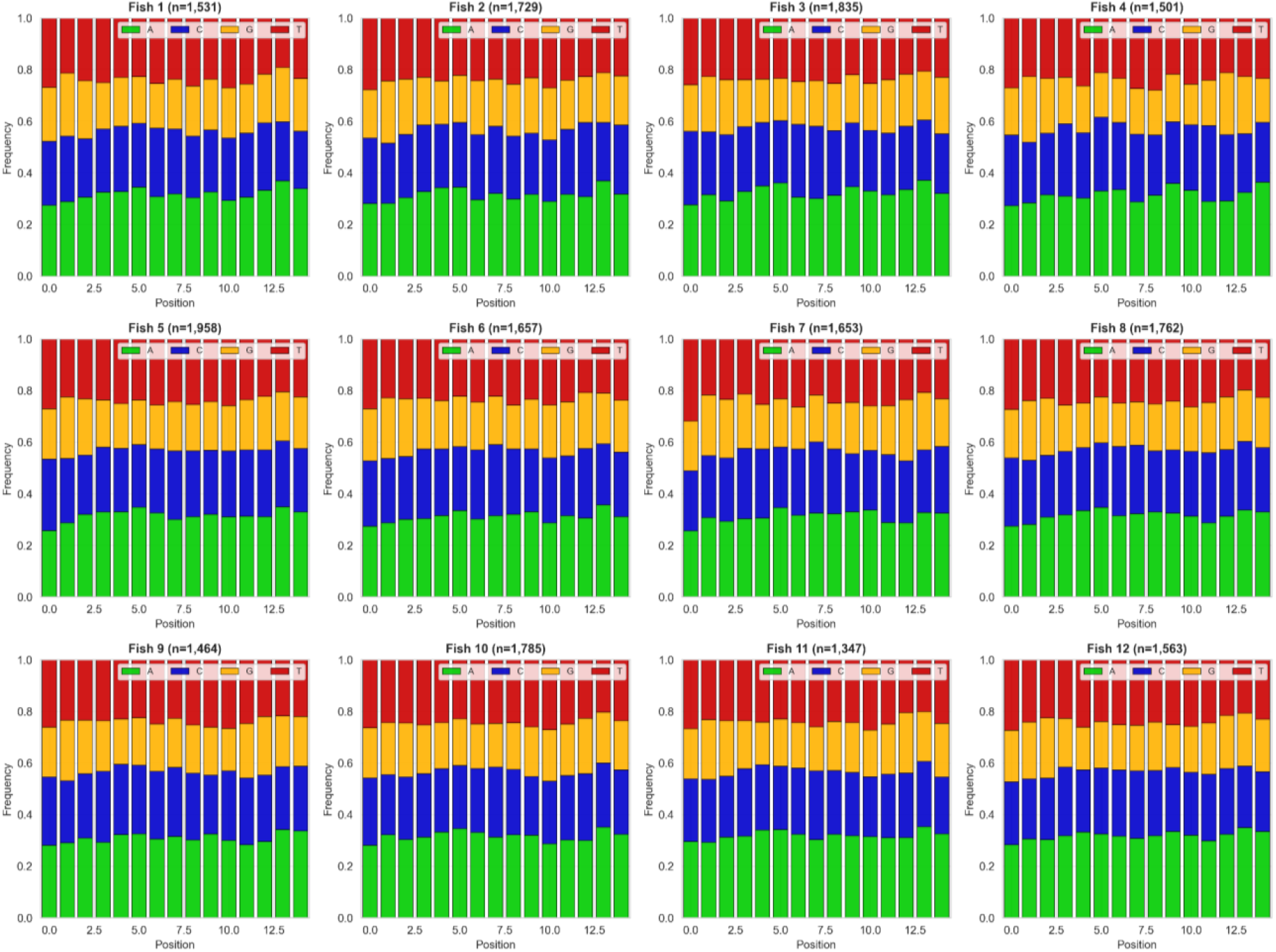
Nucleotide composition for each position in the set of barcodes recovered from each fish, related to Fig. 3. DNA sequence logos showing the positional nucleotide frequencies across all unique barcodes recovered from each of the 12 individual fish. Only 15-nt barcodes were included in the frequency calculations, although an additional minority of barcodes were 14 or 16 nt long (<1%). For each position, the height of each colored segment represents the proportion of barcodes containing that nucleotide at that position. The number in parentheses in the subtitle for each plot indicates the total number of unique barcodes analyzed for that fish sample. The close to uniform nucleotide frequencies across all positions show a similar lack of sequence bias in the recovered barcode population for all the animals analyzed.

**Figure S4.**
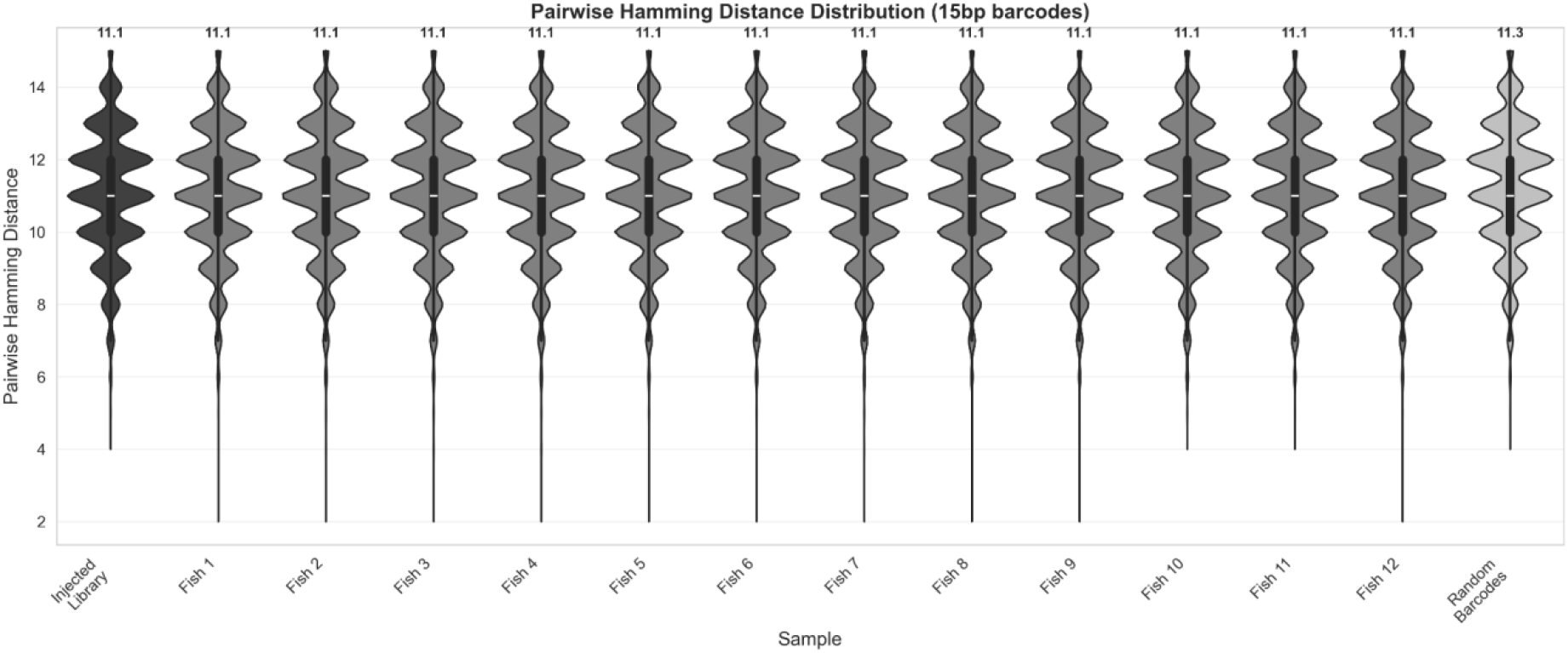
Comparison of the distribution of pairwise Hamming distances for the sets of barcodes integrated in each fish, the original injected library, and a theoretical library of uniformly distributed random 15-nt barcodes. Distributions of pairwise distances are shown for 20,000 random pairs taken from each set. On top of each violin plot there is an overlaid boxplot, showing the median and interquantile range (IQR, representing 25th and 75th percentiles), and whiskers extending to 1.5×IQR beyond the quartiles. The number above each violin plot shows the mean Hamming distance in each sample. The narrow distribution centered around Hamming distance 11.1 (close to the theoretical maximum for random sequences) indicates that barcodes are highly dissimilar to each other, confirming minimal sequence clustering or bias in the samples. Furthermore, it confirms that the integrated barcodes from the mosaic fish retained the same sequence diversity as the original injected library.

**Figure S5.**
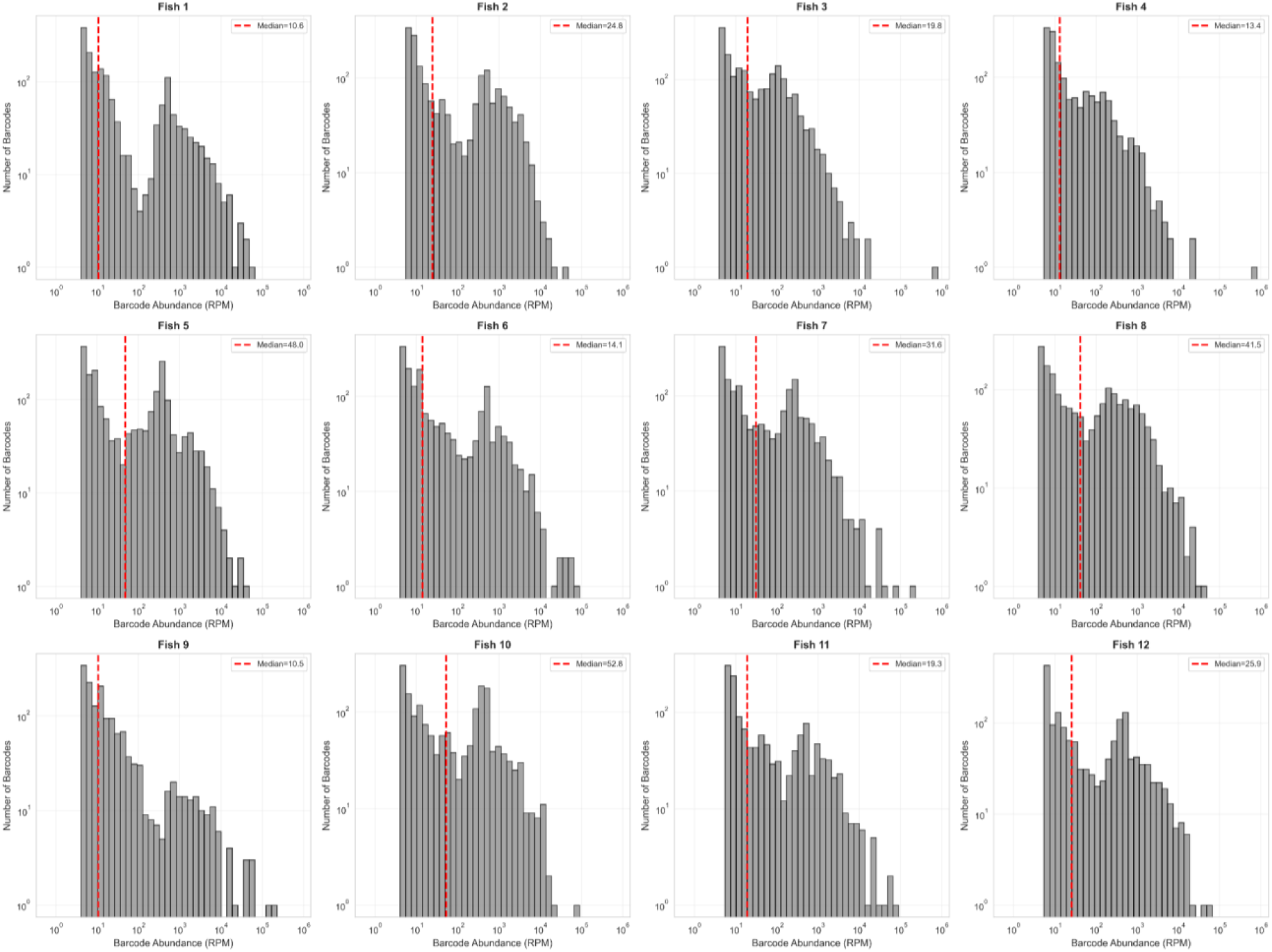
Distribution of abundance for the set of barcodes recovered from each fish, related to Figure 3. RPM=reads per million.

**Figure S6.**
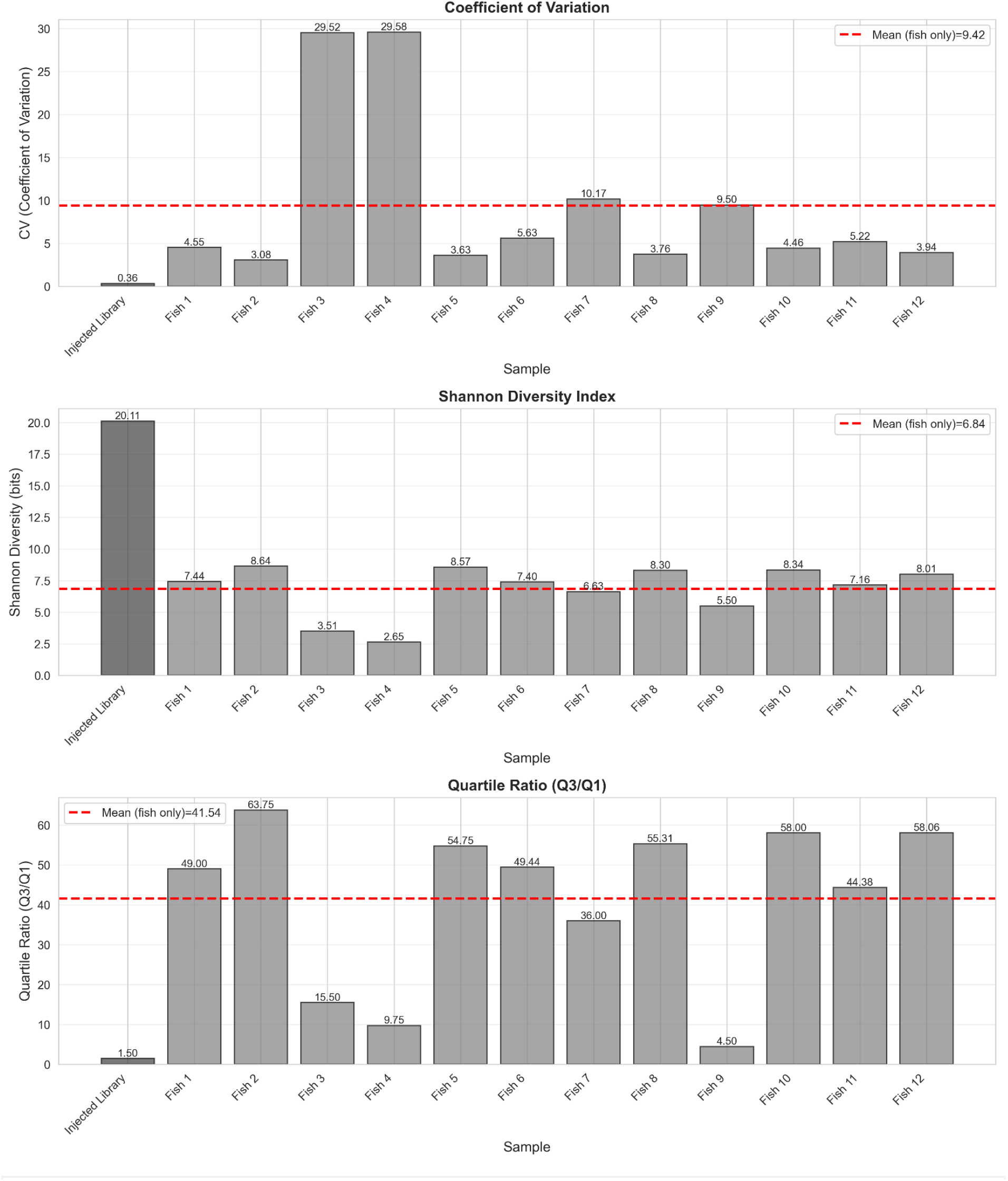
Abundance distribution metrics for the injected library and integrated barcodes recovered from each fish, quantifying the extent of clonal expansion heterogeneity and the proliferative differences among cells that received different barcode integrations. The number above each bar plot shows the value for that sample. Overall, higher CV, lower Shannon diversity and higher quartile ratios show an increase in skewness of the abundance distribution of the barcode represented in each fish, consistent with intra-fish clonal expansion of integrated barcodes. The CV (coefficient of variation) is calculated as the standard deviation divided by mean of barcode read counts. A higher CV means some barcodes are much more abundant than others (uneven distribution), while a lower CV indicates more uniform barcode representation.Shannon diversity is an entropy-based metric that considers both the number of unique barcodes and the evenness of their distribution (in bits). It quantifies the overall barcode diversity, accounting for both richness (how many different barcodes) and evenness (how uniformly distributed their abundances are). Higher values indicate more diverse, more evenly distributed barcode populations. The quartile ratio is calculated as the ratio of the 75th percentile to the 25th percentile of barcode abundance, quantifying the spread of the middle 50% of barcode abundances. Higher ratios indicate greater inequality in barcode representation.

**Table S1.**
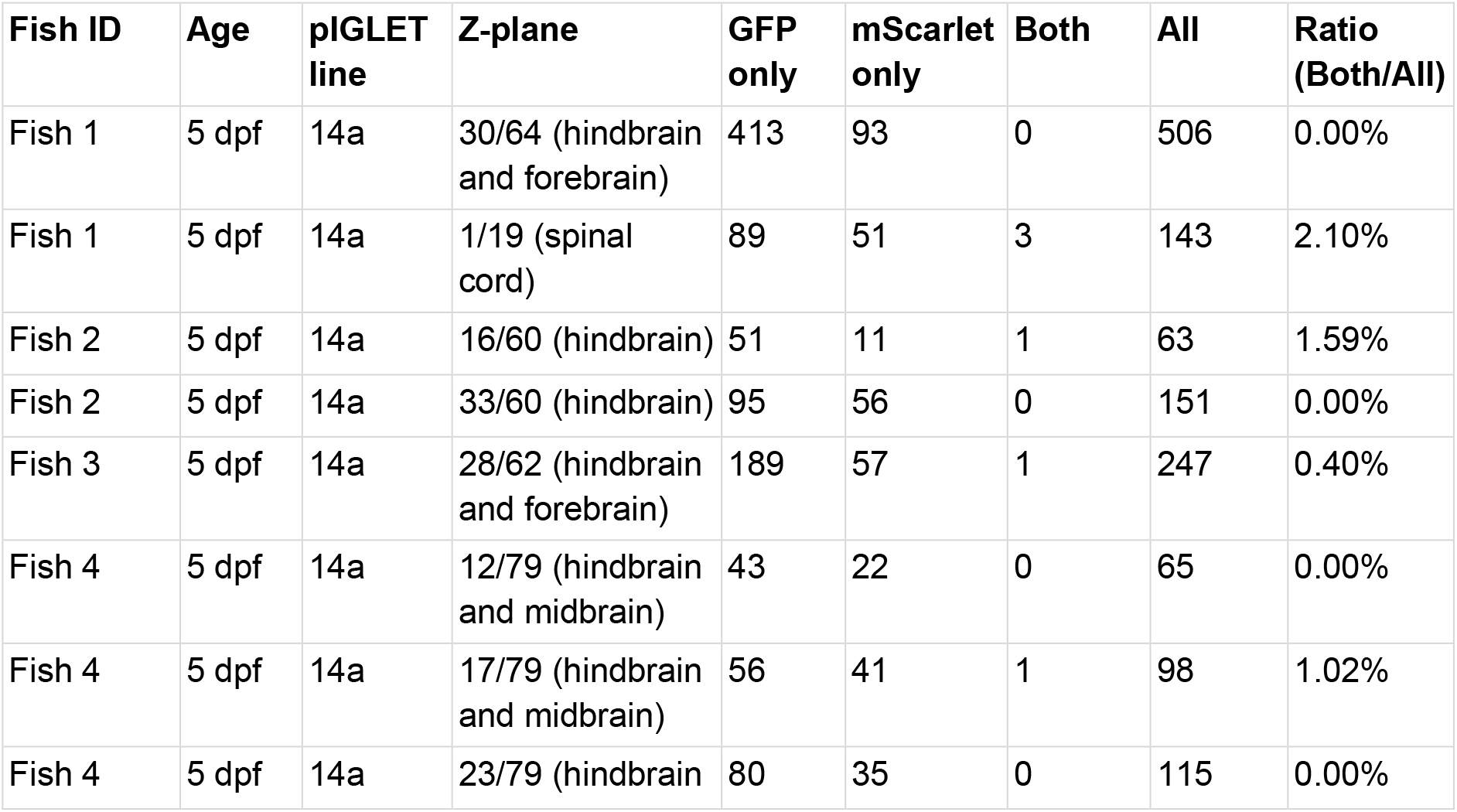
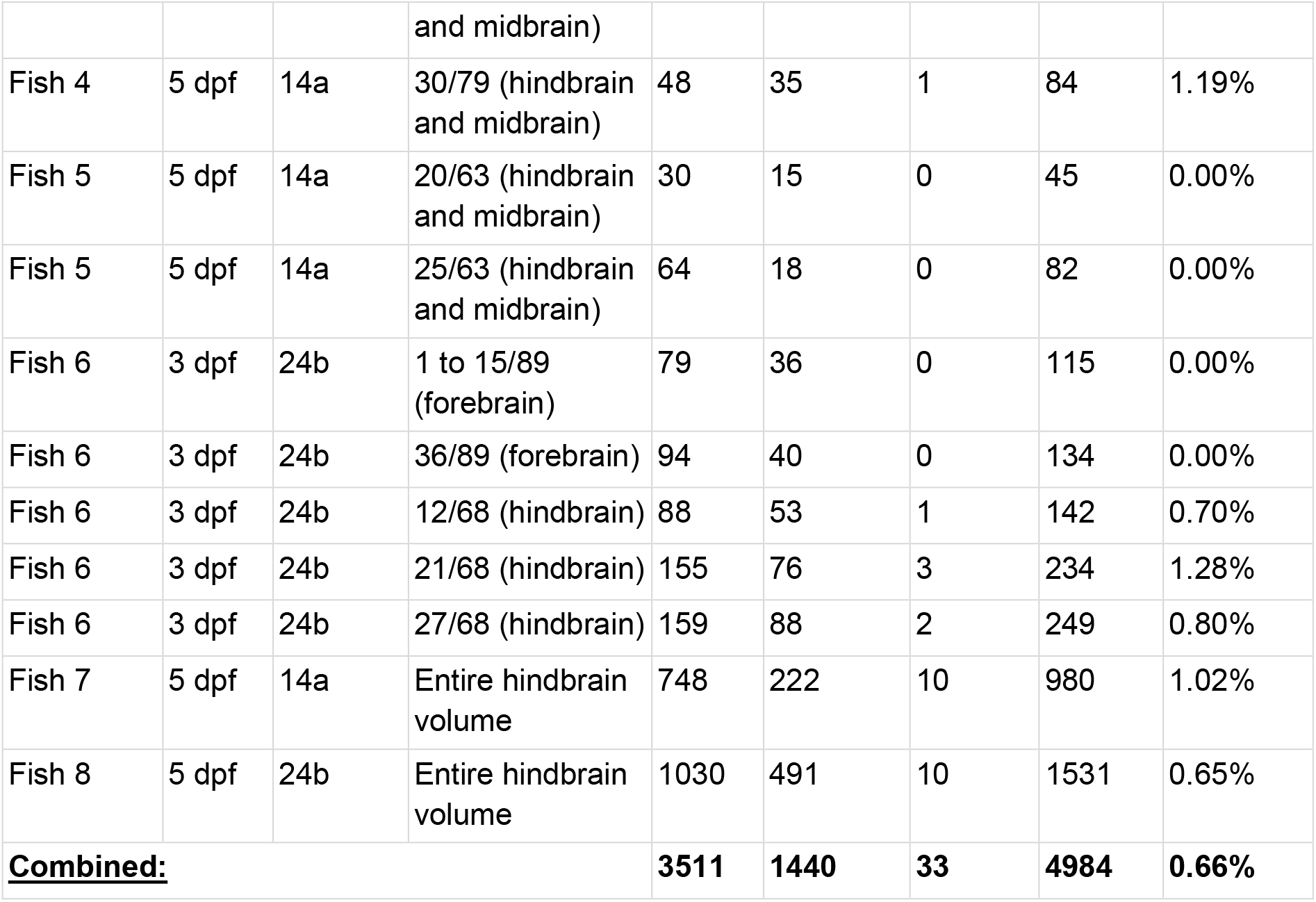
Ratio of multi-transgene neurons in the brains of mosaic zebrafish. Quantification of neurons expressing GFP-only, mScarlet-only, or both fluorescent transgenes in mosaic transgenic zebrafish brains. **Z-plane:** The specific optical section(s) analyzed from the confocal Z-stack, indicated as the plane number out of the total stack depth (e.g., “30/64” means plane 30 from a 64-plane stack). Brain region(s) visible in each plane are indicated in parentheses. For Fish 7 and 8, the entire hindbrain volume was analyzed rather than a single plane. **GFP/mScarlet only:** Number of neurons expressing only the GFP or mScarlet transgene. **Both:** Number of neurons co-expressing both the GFP and mScarlet transgenes. **All:** Total number of transgene-positive neurons counted (GFP only + mScarlet only + Both). **Ratio (Both/All):** Percentage of all transgene-positive neurons that express both fluorophores, compared to all counted neurons. This ratio is used to estimate the frequency of multi-transgene integration events. **Combined**: Summary statistics pooling all the analyzed planes across all fish.

**Table S2.** Sequencing library quality metrics. Quality metrics for the Illumina sequencing libraries from the injected barcode library and pooled integrated barcodes extracted from 12 individual fish. **Mean quality score**: Average Phred quality score across all base calls in the library. The Phred quality score is a logarithmic measure of base calling accuracy, calculated as Q = -10 × log_10_(P), where P is the probability of an incorrect base call. Q20: 1% error rate (99% accuracy), Q30: 0.1% error rate (99.9% accuracy). **Bases ≥ Q20**: Percentage of sequenced bases with Phred quality score of 20 or higher. **Bases ≥ Q30**: Percentage of sequenced bases with Phred quality score of 30 or higher.

|  | **Injected barcode library** | **Integrated barcodes (12 pooled fish samples)** |
| --- | --- | --- |
| **Sequenced reads** | 5,822,820 | 7,191,895 |
| **Mean quality score:** | 39.13 | 39.22 |
| **Bases ≥ Q20:** | 99.13% | 99.02% |
| **Bases ≥ Q30:** | 95.24% | 95.84% |

**Table S3.**
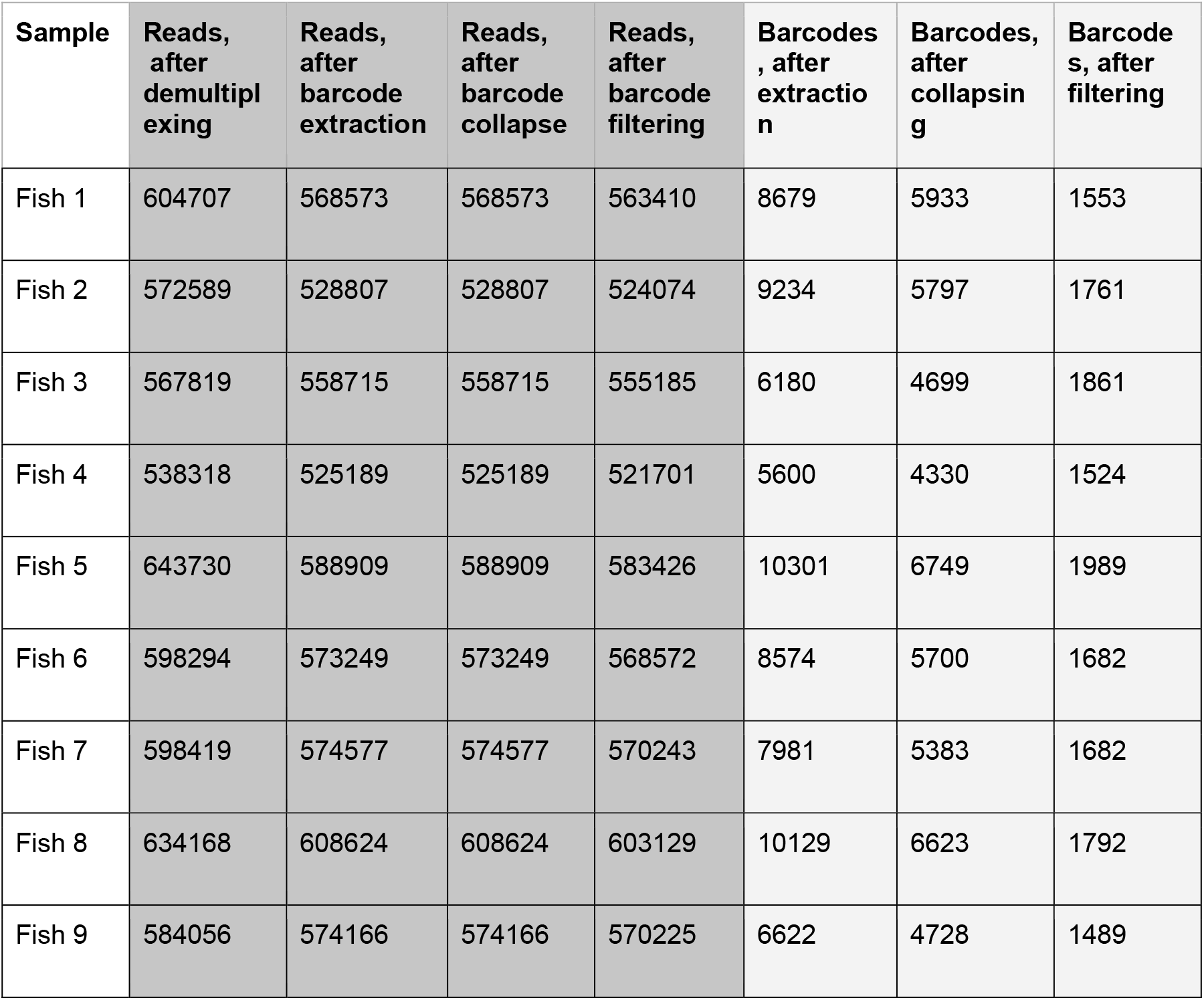
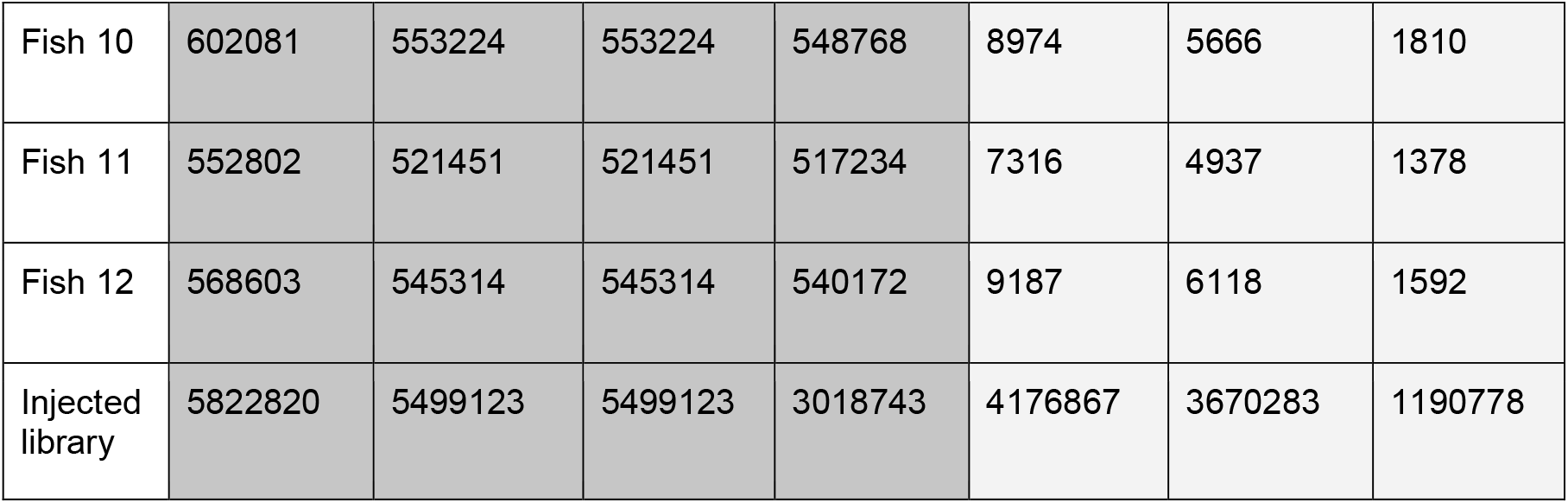
barcode counts and read retention throughout the stages of barcode extraction and processing. Reads, after demultiplexing: Total number of paired-end sequencing reads assigned to each sample. For fish samples, reads were demultiplexed based on 5bp sample barcodes at the read start, allowing up to 1 mismatch for error correction. The injected library was sequenced separately and required no demultiplexing. **Reads, after barcode extraction:** Number of reads from which valid barcodes were successfully extracted. Extraction required identifying conserved anchor sequences flanking the random barcode region. Reads lacking proper anchor sequences or with barcodes outside expected positions and lengths (14-16 nt) were discarded. **Reads, after barcode collapse:** Number of reads remaining after merging similar barcodes. Collapsing merges counts into parent barcodes but does not discard reads. **Reads, after barcode filtering:** Number of reads associated with barcodes that passed abundance and sequence filters. For the fish samples, barcodes were excluded from a fish if they appeared <3 times in that fish. For the injected library, barcodes with <2 reads in the injected library were excluded. Additionally, barcodes too similar (Levenshtein distance ≤2) to the conserved non-barcode regions of the injected plasmid were removed. **Barcodes, after extraction:** Number of unique barcode sequences identified after extraction, before any quality filtering or collapsing. **Barcodes, after collapsing:** Number of unique barcode sequences after collapsing. Barcodes within Levenshtein distance of 1, assumed to result from PCR or sequencing errors, were merged into their most abundant neighbor (parent barcode), with read counts combined. **Barcodes, after filtering:** Number of unique barcodes retained after applying the abundance and sequence filters described above.

**Appendix data S1:**
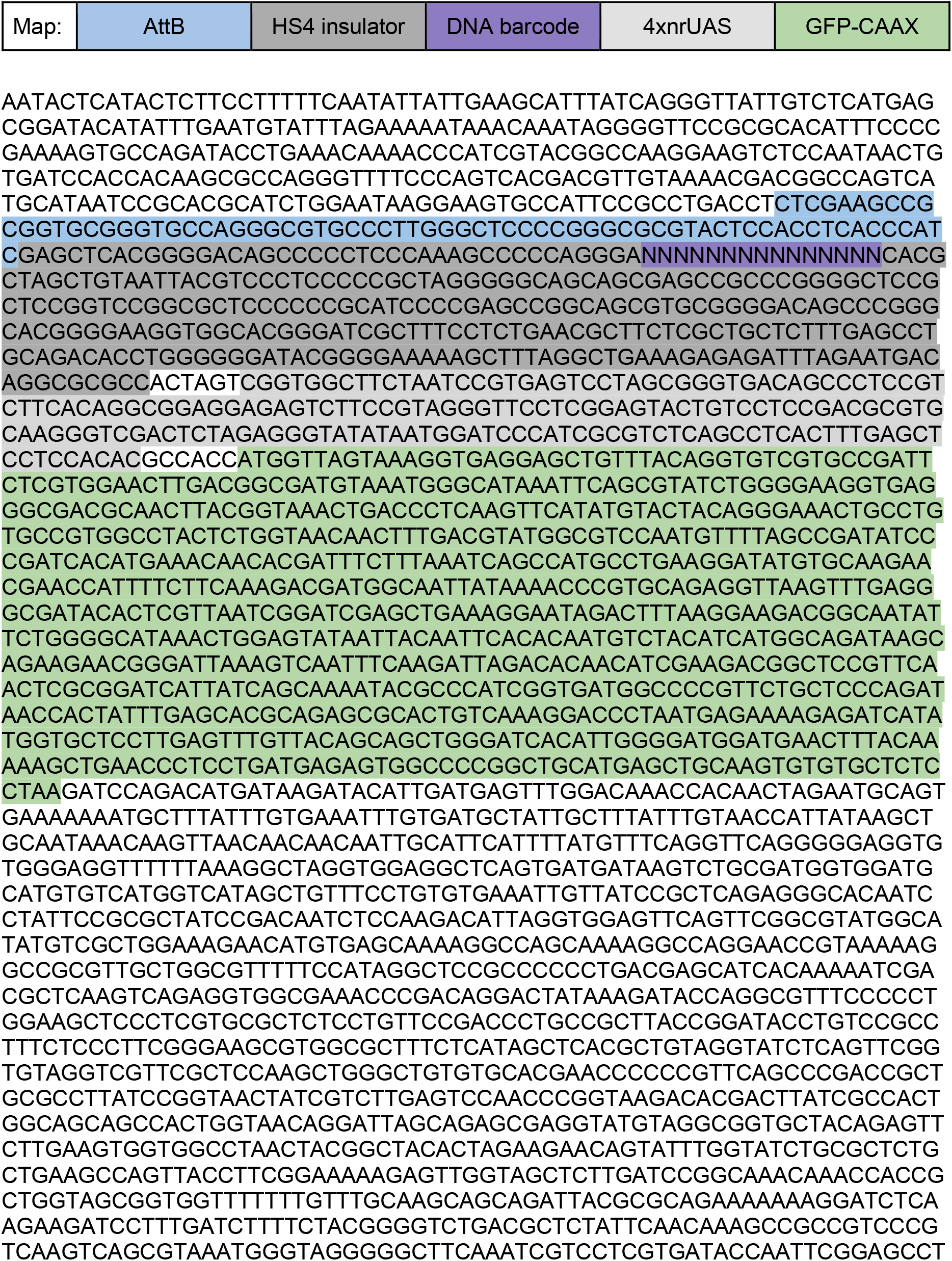
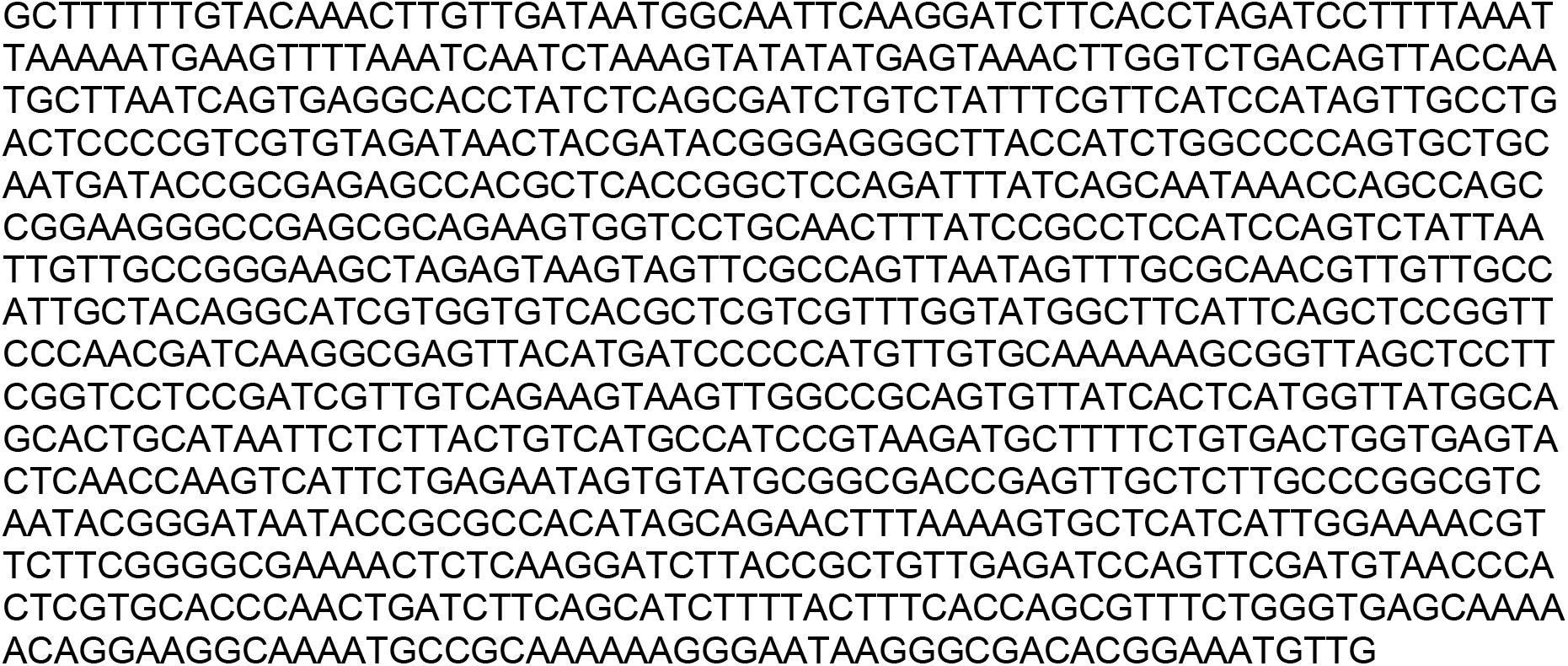
barcoded AttB-HS4-15N-nrUAS-GFP-CAAX plasmid sequence.

**Appendix data S2:**
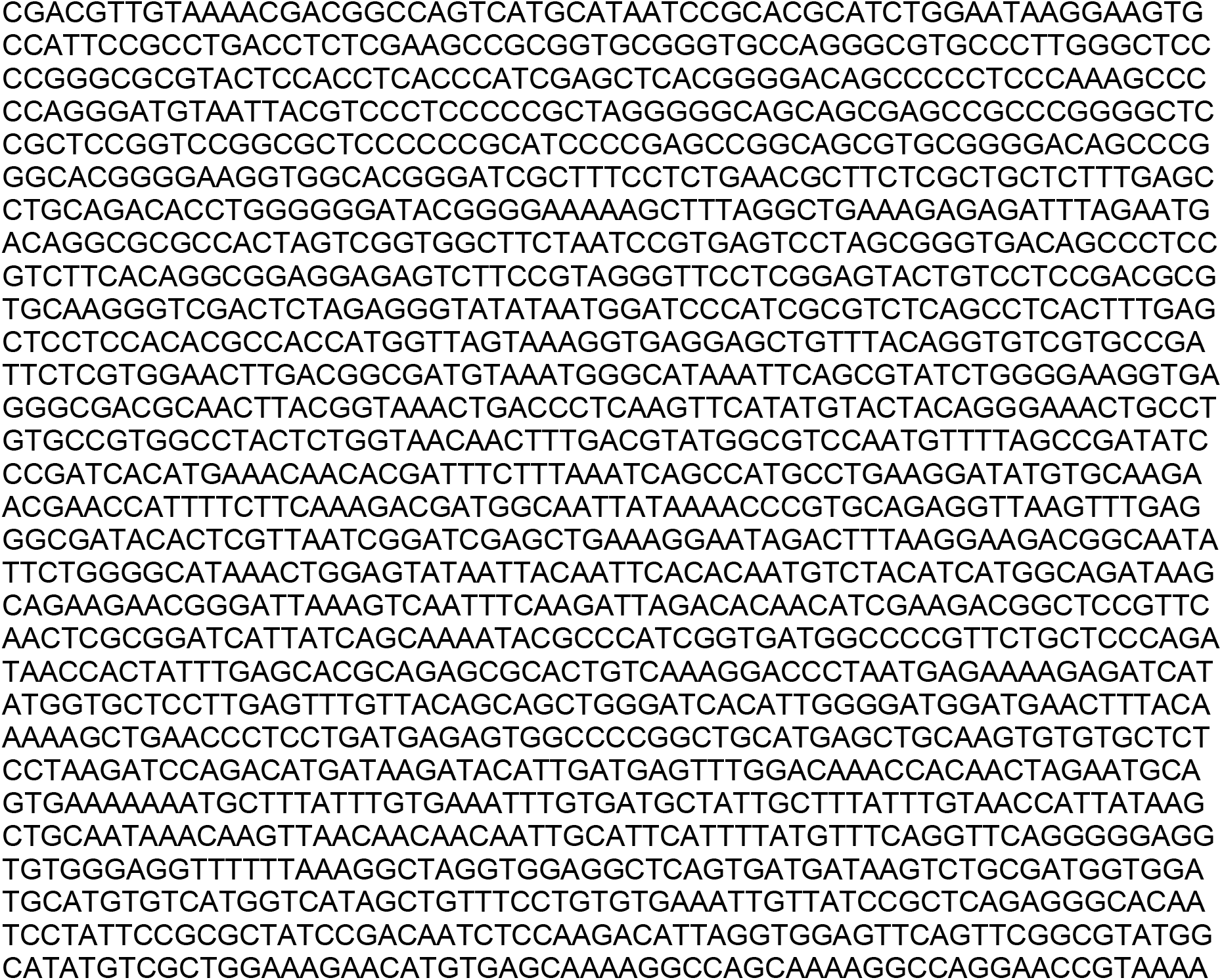
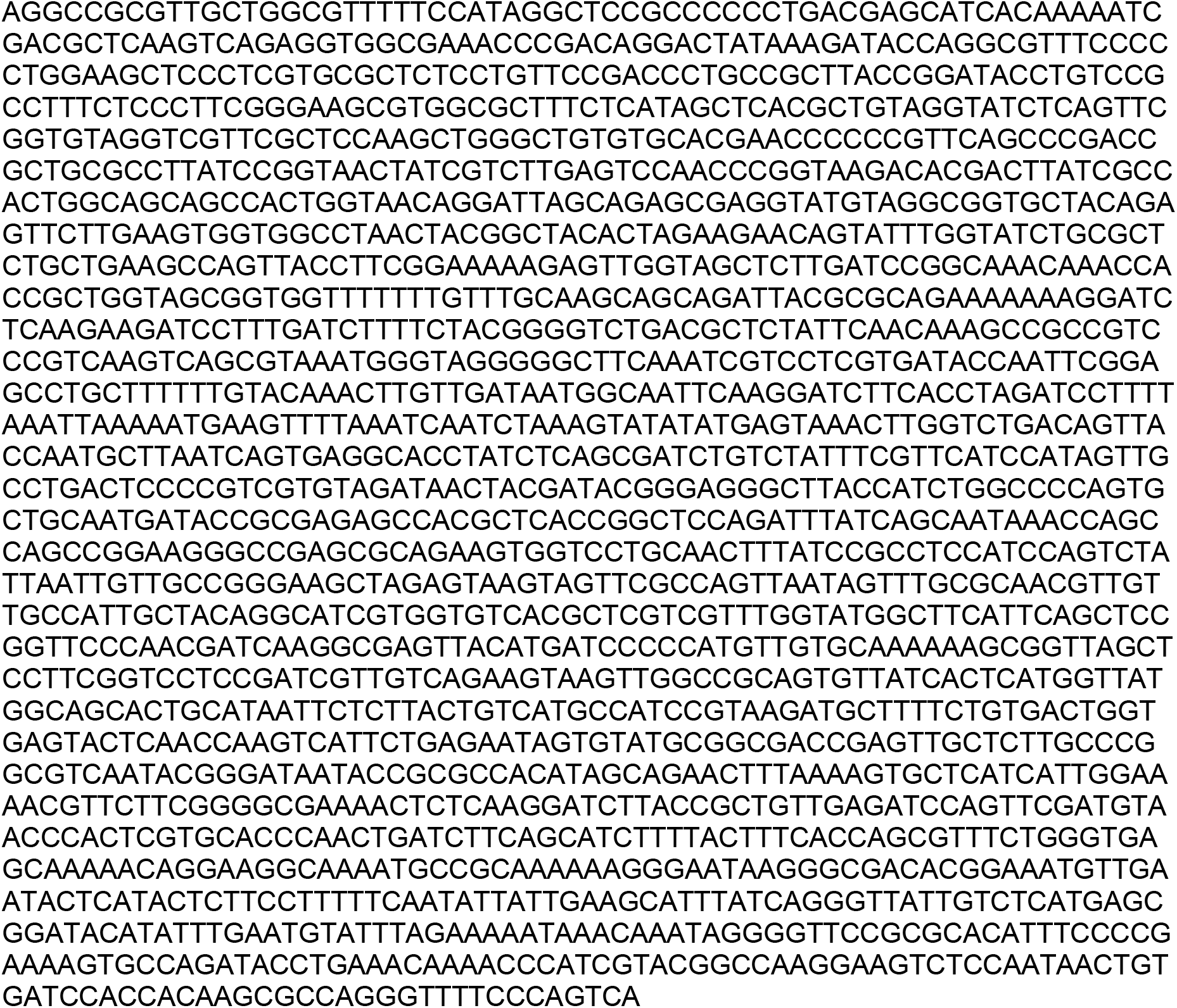
AttB-HS4-nrUAS-GFP-CAAX plasmid sequence.

**Appendix data S3:**
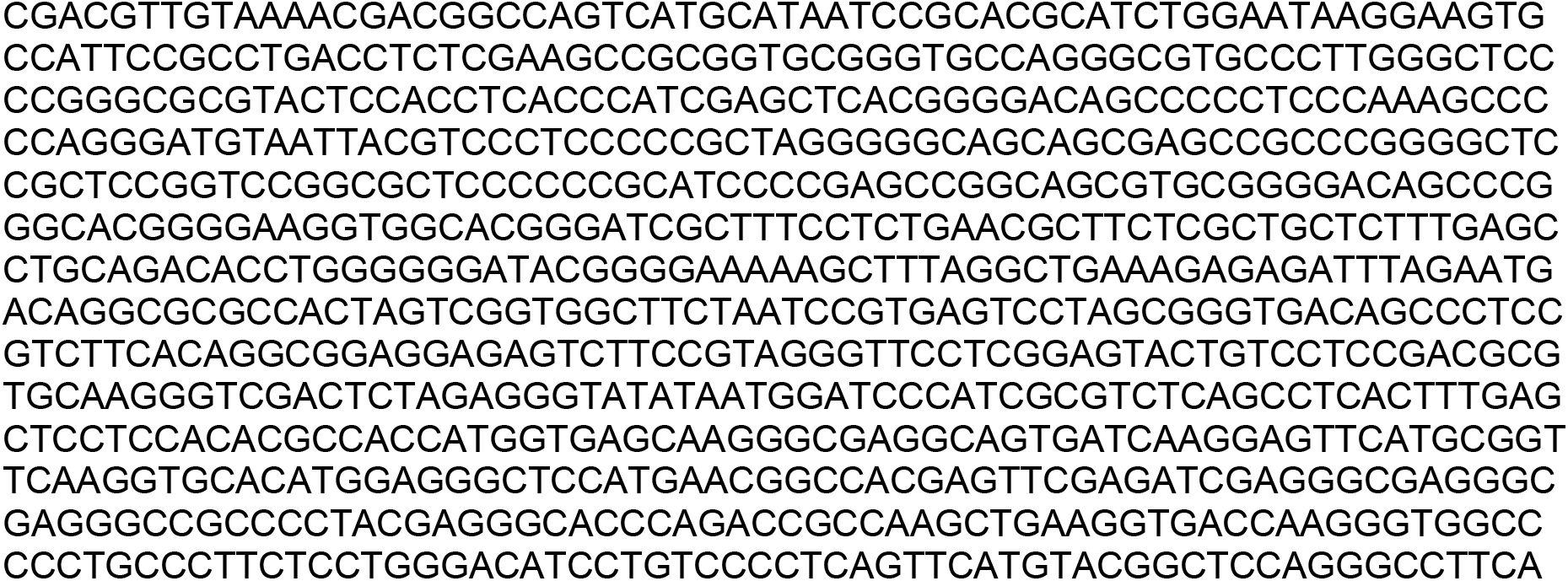
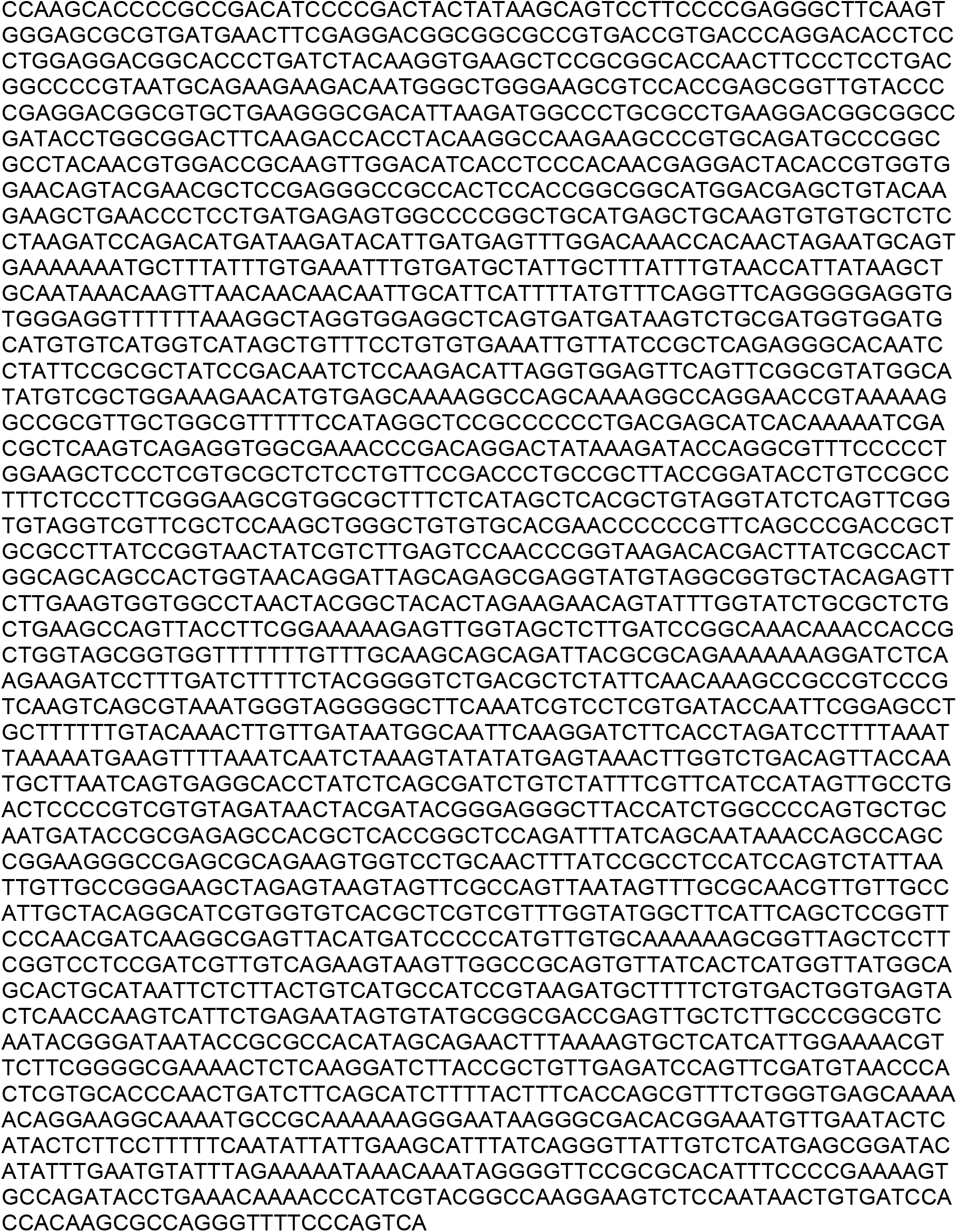
AttB-HS4-nrUAS-mScarlet-CAAX plasmid sequence.

**Appendix data S4:**
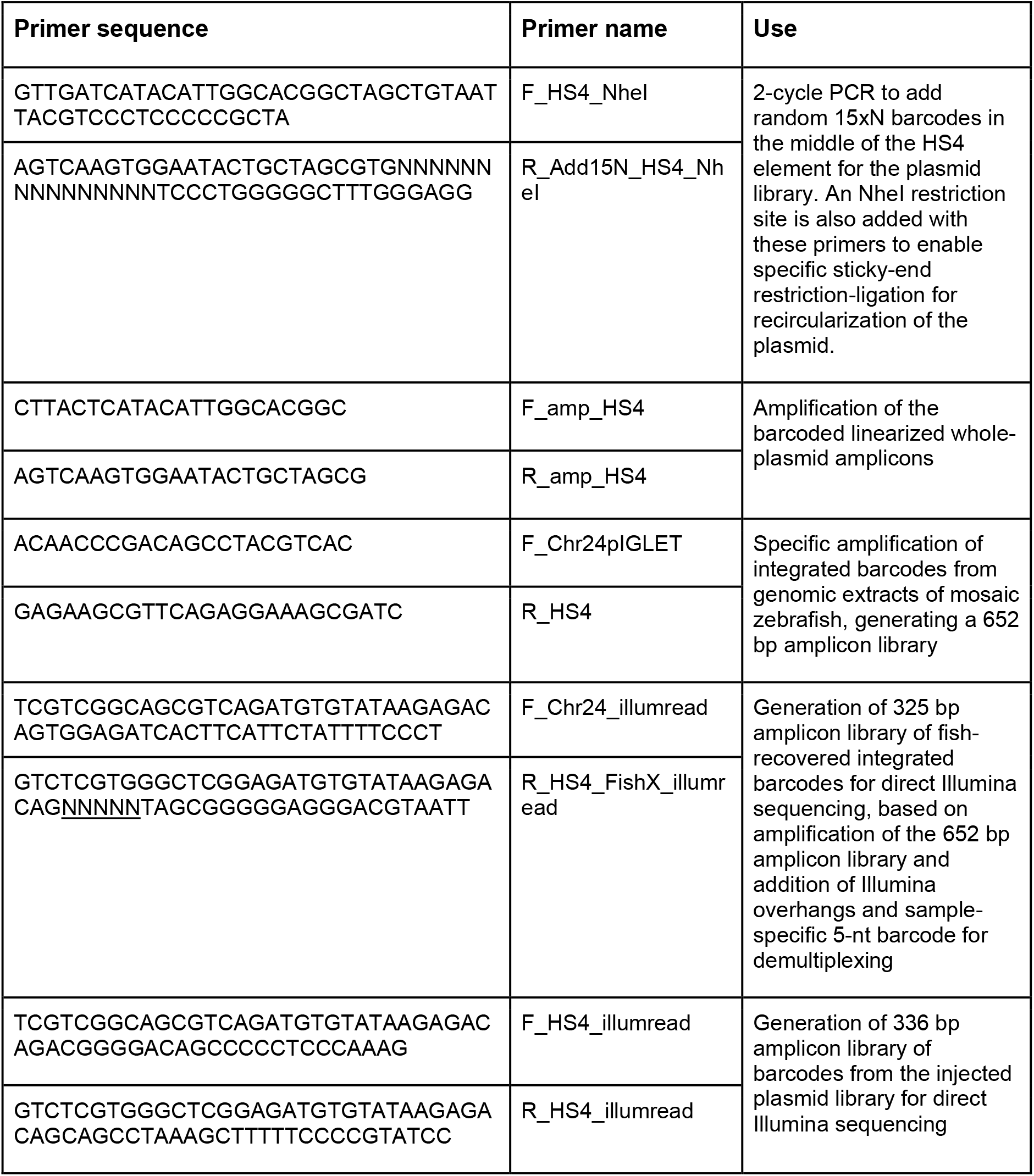
Sequences of key primers used for library generation and sequencing.

## Additional information

### Contact information

Edward Boyden

### Data sharing plans

All the raw imaging data and Illumina sequencing data associated with Fig. 2, 3, S1, S2, S3, S4, S5, S6, and Table S1, S2, S3, are available on DOI https://doi.org/10.5061/dryad.d2547d8h0.

All the code used to analyze the sequencing data is available on https://github.com/shaharbr/library_transgenesis. The full sequences of key plasmids and primers used in this study are available in appendix data S1, S2, S3 and S4.

### Significance statement

Genetic perturbations and molecular tools characterized in cell culture frequently fail to translate in vivo, yet pooled screening in living animals faces critical limitations: the high prevalence of multi-transgene cells confounds interpretation, viral packaging constrains transgene size, and tropism introduces biases. We developed a library transgenesis method, implemented in zebrafish, that overcomes these challenges by exploiting delayed site-specific integration to create mosaic animals with >1,500 multi-kilobase transgenes integrated per animal. In those library mosaics, ∼99% of the cells express a single library member, thanks to the mutual-exclusivity enforced by the site-specific integration mechanism. Library transgenesis can transform each animal into hundreds of parallel experiments, enabling direct in vivo screening of molecular tools and genetic perturbations in their native physiological contexts.

